# ICAT: A Novel Algorithm to Robustly Identify Cell States Following Perturbations in Single Cell Transcriptomes

**DOI:** 10.1101/2022.05.26.493603

**Authors:** Dakota Y. Hawkins, Daniel T. Zuch, James Huth, Nahomie Rodriguez-Sastre, Kelley R. McCutcheon, Abigail Glick, Alexandra T. Lion, Christopher F. Thomas, Abigail E. Descoteaux, W. Evan Johnson, Cynthia A. Bradham

## Abstract

**Motivation:** The detection of distinct cellular identities is central to the analysis of single-cell RNA sequencing experiments. However, in perturbation experiments, current methods typically fail to correctly match cell states between conditions or erroneously remove population substructure. Here we present the novel, unsupervised algorithm ICAT that employs self-supervised feature weighting and control-guided clustering to accurately resolve cell states across heterogeneous conditions.

**Results:** Using simulated and real datasets, we show ICAT is superior in identifying and resolving cell states compared to current integration workflows. While requiring no a priori knowledge of extant cell states or discriminatory marker genes, ICAT is robust to low signal strength, high perturbation severity, and disparate cell type proportions. We empirically validate ICAT in a developmental model and find that only ICAT identifies a perturbation-unique cellular response. Taken together, our results demonstrate that ICAT offers a significant improvement in defining cellular responses to perturbation in single-cell RNA sequencing data.

**Availability and implementation:** https://github.com/BradhamLab/icat

Supplemental Methods, Tables and Figures are available online.

## Introduction

From deconstructing tumor cell compositions (Schelker *et al*., 2017), to reconstructing developmental pathways (Wagner *et al*., 2018), single-cell RNA sequencing (scRNA-seq) has revolutionized scientists’ ability to explore complex tissues and systems. Clustering of individual cells is central to the analysis of scRNA-seq data, in which cell states are identified by grouping cells with similar gene expression profiles (Luecken and Theis, 2019). Whether to ascertain the effect of drug treatment on cancer viability or to infer the mechanistic role of a given gene in tissue specification, perturbation experiments have long been used to dissect and understand complex biological systems. Oftentimes this analysis is formalized by identifying changes in the abundance of cell states between treatments (Luecken and Theis, 2019; Kang *et al*., 2017; Haber *et al*., 2017). Inherently, such analyses are based on the assumption that cell states are readily identified and matched between treatments. Recently, graph-based community detection algorithms such as the Louvain and Leiden methods have become best practice for identifying cell states in single-condition scRNA-seq datasets (Luecken and Theis, 2019). However, these methods often fail to match identities in datasets containing perturbing treatments, which present particular difficulties due to dominating treatment effects (Fig. S1), where it is typical that scRNA-seq datasets segregate by treatments rather than cellular identities (Fig. S1), similar to the impact of batch effects (Luecken and Theis, 2019; Kang *et al*., 2017; Lähnemann *et al*., 2020; Kagohara *et al*., 2020). This challenge presents significant difficulties in correctly evaluating cellular responses to perturbation in scRNA-seq experiments.

Common techniques for identifying cell states in scRNA-seq perturbation experiments rely on either reference datasets to map cells onto known cell states (Regev *et al*., 2017; Schaum *et al*., 2018), or an integration algorithm to minimize differences between control and perturbed cells by transforming cells into a shared space prior to clustering cells (Luecken and Theis, 2019; Perillo *et al*., 2020). The first scenario is problematic for non-model systems, where such atlases may not exist, and is inappropriate when the goal of a given experiment is discovery of novel cell states or subtypes; for these reasons, the latter approach is often preferred. Toward that end, scRNA-seq algorithms have been developed that broadly integrate heterogeneous datasets (Luecken and Theis, 2019; Stuart and Satija, 2019; Hie *et al*., 2019). Originally developed to remove batch effects (Haghverdi *et al*., 2018), many of the most widely used integration algorithms work by matching mutual nearest neighbors between batches in order to learn non-linear transformations that minimize neighbor distance in an embedded space (Haghverdi *et al*., 2018; Stuart and Satija, 2019; Hie *et al*., 2019; Korsunsky *et al*., 2019).While useful for removing technical variation from datasets, integration algorithms are unable to discriminate technical from biological noise: thus their use may be inappropriate when biological differences exist between samples, such that a one-to-one correspondence between cell states cannot be assumed. This difficulty is reinforced by recent findings which show that integration methods often smooth over and eliminate biologically meaningful signals, leading to incorrect cell state identification and possibly wholesale erasure of real cell states (Luecken and Theis, 2019; Luecken *et al*., 2022; Büttner *et al*., 2019; Tyler *et al*., 2021).

We therefore developed a two-step algorithm to robustly and sensitively **<u>I</u>**dentify **<u>C</u>**ell states **<u>A</u>**cross **<u>T</u>**reatments (ICAT) in scRNA-seq data. In contrast to integration approaches that minimize differences between samples, ICAT uses a novel approach that relies on self-supervised feature selection and control-guided clustering to identify cell states using biologically relevant features. While other methods exist to either identify shared clusters between treatments (Barron *et al*., 2018) or to leverage sparse features (Hua *et al*., 2020), to our knowledge, ICAT is the only method capable of doing both while also handling multiple experimental conditions. Our results show that, by emphasizing cell-state defining genes, ICAT reliably identifies cell states across scRNA-seq perturbation experiments with high accuracy. Importantly, ICAT does not require prior knowledge of marker genes or extant cell states, is robust to perturbation severity, identifies cell states with higher accuracy than leading integration workflows with both simulated and real scRNA-seq perturbation experiments, and operates reliably at low signal resolutions.

## Materials and Methods

### ICAT Algorithm overview

To overcome the inability of current clustering algorithms to accurately detect cell states in perturbation experiments (Fig. S1), we developed a two-step algorithm (Fig. 1A). The purpose of the first step is to determine the cluster-defining features and weight them among the control cells alone. The goal of the second step is to use those feature weights to co-cluster the controls and the perturbed cells together, effectively assigning the perturbed cells to control clusters. Perturbed cells may also be optionally clustered separately to determine their feature weights in order to detect cell states that are asymmetrically present in the perturbed condition. Using the Louvain method (Blondel *et al*., 2008) by default, the first step separately clusters control cells to produce initial cluster labels for each cell. Cluster labels are then used to weight genes by their ability to predict the previously generated labels using Neighborhood Component Feature Selection (NCFS) (Yang *et al*., 2012). NCFS weights genes by using gradient ascent to maximize a regularized leave-one-out classification accuracy metric. By the nature of regularization, most gene weights converge to zero, leaving only the most predictive genes with weights greater than 1. Applying the learned gene weights to the original expression matrix with both control and treated cells transforms the matrix into a “cluster-defined” space, where the distances between cells are dominated by highly weighted marker genes. After this transformation, ICAT retains previously identified cluster labels for control cells, then, in the second step, performs semi-supervised Louvain community detection (Blondel *et al*., 2008; Traag, 2021), whereby control labels are held immutable (Fig. 1A).

**Figure 1.**
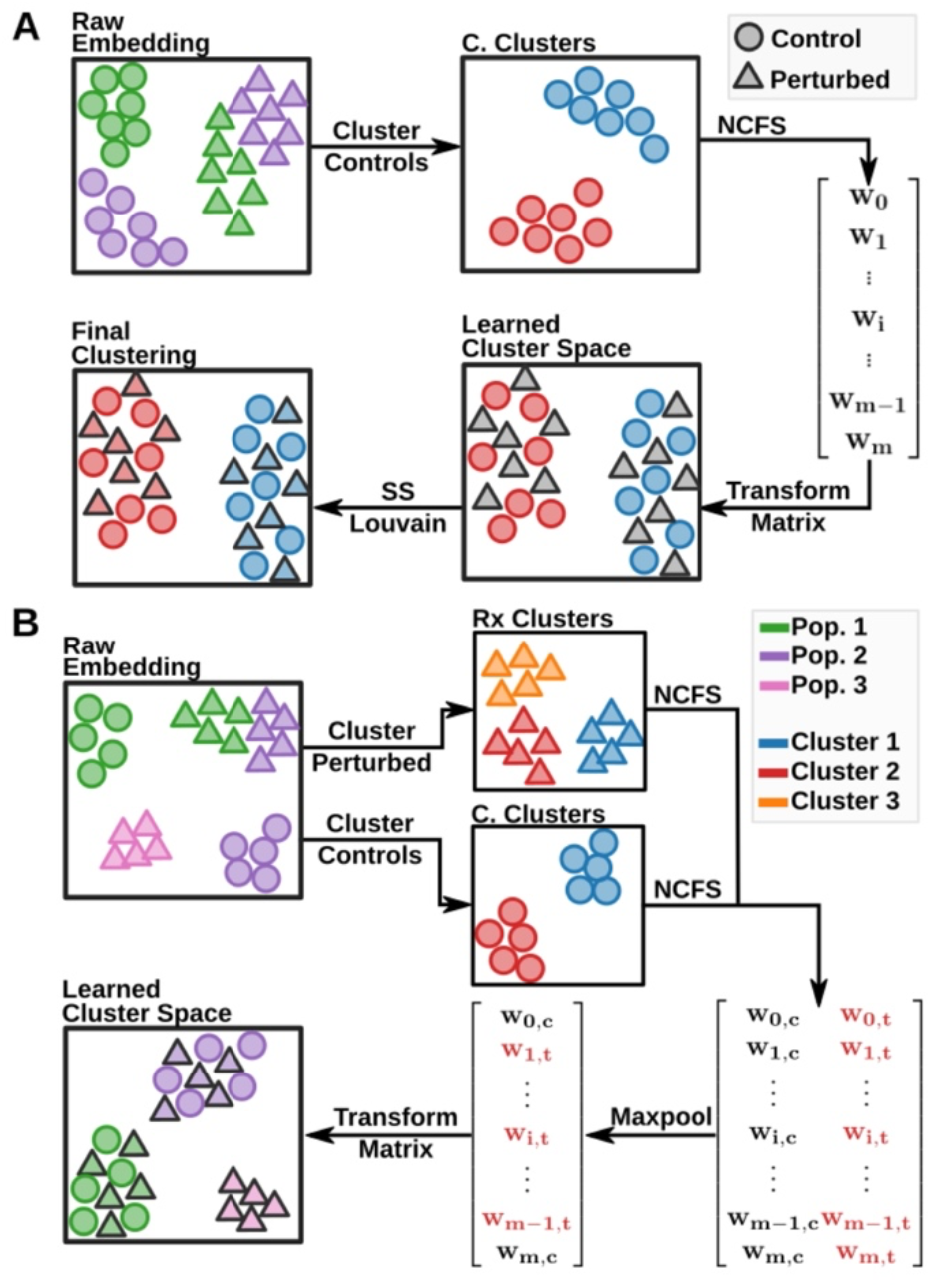
Overview of the ICAT algorithm. **A**. The schematic illustrates the ICAT_C_ implementation of ICAT. To identify cell states across treatments, ICAT first performs self-supervised feature weighting to find genes that discriminate cell identities among control cells alone, followed by semi-supervised clustering using the newly transformed expression matrix. To learn feature weights, ICAT clusters control cells using Louvain community detection, then uses the newly generated cluster labels as input into NCFS to weight genes by their predictiveness. After applying the learned gene weights to the original gene expression matrix, ICAT clusters both treated and control cells together using a semi-supervised implementation of Louvain community detection. During this process, ICAT holds the previously identified cluster labels for the control cells immutable. **B**. The schematic illustrates the ICAT_C+T_ implementation, which expands feature weighting to treated cells to identify asymmetrical populations between treatments. Cells are split along treatments and independently clustered using the Louvain method, then cluster labels are used to learn gene weights using NCFS in each treatment set independently. To retain asymmetrically informative genes, weights for each treatment are concatenated row-wise and subsequently reduced to the maximum weight using a row-wise maxpool function. The reduced weight vector is then used to transform the original count matrix.

This strategy achieves three goals: 1. Because NCFS is a sparse method, it overtly separates informative and non-informative genes, making it possible to cluster directly on cell-state defining genes while simultaneously removing likely sources of biological noise. 2. ICAT produces easily interpretable results by clustering on the expression patterns of a few, highly weighted genes. This is a noted benefit over typical integration methods, where transformations are obscured behind neighbor-based approaches (Stuart and Satija, 2019; Hie et al., 2019). 3. ICAT makes comparing treatment effects on cell states straightforward. By implementing semi-supervised clustering with immutable control labels, ICAT ensures cell states discovered in control cells are retained when treated cells are introduced, alleviating concerns of artifact-induced cell states upon integration due to erroneously grouping cells. These three features together produce interpretable results that ease compositional analysis between treatments.

While similar in approach, ICAT provides several advantages compared to identifying shared cell states via marker gene mapping: first, ICAT does not require previously known marker genes, making ICAT appropriate for situations with previously uncharacterized cell states. Second, by clustering control and treated cells together, ICAT takes advantage of the structure in both datasets compared to clustering both independently. This alleviates issues with imperfect matches between marker genes. Third, by foregoing common marker gene presence-absence comparisons to map cell types, ICAT accurately identifies activated states, which is otherwise difficult. Finally, by generating cell state labels through a clustering step rather than a classification approach, ICAT is able to group cells with treatment-specific phenotypes and identify out-of-sample states.

One caveat of our approach is that if asymmetrical populations exist in treated cells, but not controls, their identifying genes will likely be collapsed by the control-only feature-weighting step. Depending on the preferred cell-state resolution, this may or may not be desirable. To account for both resolution levels, researchers can choose to perform feature-weighting in all treatment types to identify asymmetrical populations in each condition (Fig. 1B, Fig. 2A, Fig. 3A). To reduce gene weights to a single gene-weight vector, only the maximum gene weight among the conditions is retained. During semi-supervised clustering, labels for control cells are retained during label initialization, while treated cells are re-initialized as singleton clusters.

**Figure 2:**
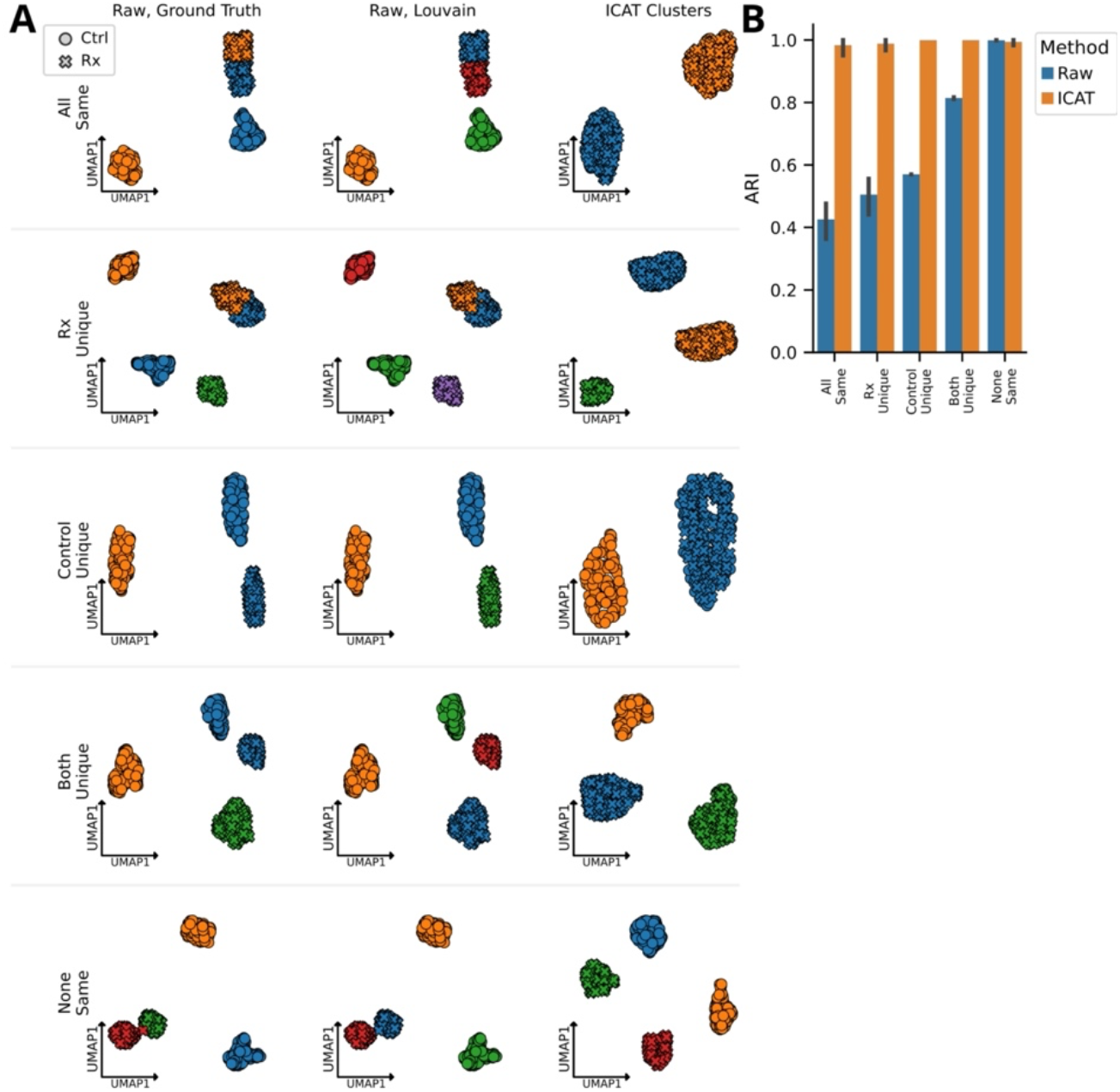
ICAT correctly identifies cell states in distinct experimental compositions. **A**. UMAP projections of different cellular compositions in simulated datasets. Each dot represents a cell with circles representing control cells and crosses denoting treated cells. Dots are colored by ground truth identity (left column), cluster label produced by performing Louvain community detection on the raw count matrix (middle column), and clusters labeled produced by ICAT (right column). **B**. Average agreement between ground truth label and cluster labels produced by clustering the raw data (blue) and ICAT (orange) as measured by the Adjusted Rand Index (ARI). Error bars represent the 95% confidence intervals for the mean ARI for each method. Five different cellular composition conditions were simulated: “All same”, both control and treated cells share the same two cell states; “Rx Unique”, treated cells contain a treatment-unique cell state; “Control Unique”, control cells contain a unique cell state; “Both Unique”, both treated and control cells contain treatment-specific cell states; and “None Same”, no shared cell states between treated and control cells. Each condition was simulated fifteen times (n = 15). Simulations were evaluated using the ICAT_C_ implementation.

**Figure 3.**
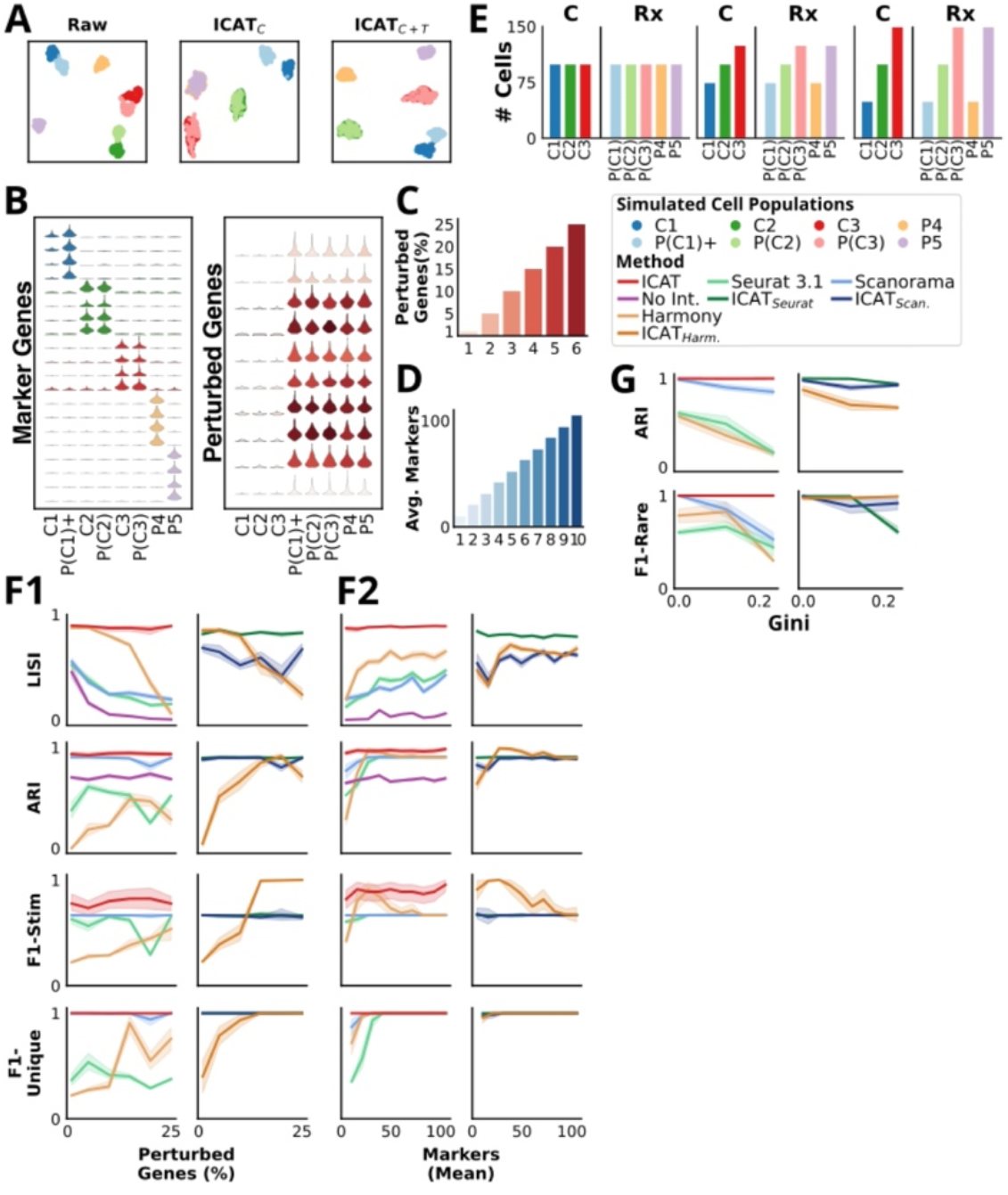
ICAT outperforms current methods for cell state identification and is robust to experimental conditions. **A**. UMAP projections of raw (left), ICAT_C_ processed (middle), and ICAT_C+T_ processed count matrices for the simulated data. Projections show ICAT_C_ and ICAT_C+T_ correctly mix shared populations (red and green), while only ICAT_C+T_ isolates asymmetrical populations (purple, yellow). ICAT performance for simulated data was further evaluated using the ICAT_C+T_ implementation only. **B**. Select marker and perturbed gene expression patterns are displayed as violin plots for three simulated control cell types (1-3) under normal (C) and perturbed (P) conditions. P(C1)+ is a stimulated and perturbed version of cell type 1; perturbation-specific cell types P4 and P5 express distinct marker genes. **C**. The percentage of perturbed genes used to assess robustness to perturbation severity is shown; values range from 1-25%. **D**. The average number of marker genes per cell identity used to assess robustness to signal strength is shown; values range from 10 to 105 (0.7 - 7% of total genes). **E**. The set of cell identity proportions used to test the ability to identify rare cell states is shown. The number of cells per treatment-label pair ranges from 50 to 175. **F**. Method performance is compared as the fraction of perturbed genes increases (F1), and as the average number of marker genes per population increases (F2). Left panels display results for stand-alone methods: ICAT, No Int. Seurat, Scanorama, and Harmony; while right panels show results for ICAT-extended workflows: ICAT_Seurat_, ICAT_Scan_, and ICAT_Harm_. Results are depicted as averages with 95% confidence intervals shown by shading. **G**. Method performance is compared as the proportion of cell types is varied. The Gini coefficient reflects the degree of population size disproportion among cell states. Left panel: ICAT, Seurat, Scanorama, and Harmony. Right panel: ICAT_Seurat_, ICAT_Scan_, and ICAT_Harm_.

### Data simulation

To assess ICAT, we initially tested performance in simulated single-cell RNAseq data (Fig. 2A, 3A-E). We simulated data using a zero-inflated Negative Binomial distribution to model gene counts per Büttner and colleagues (Büttner *et al*., 2019, see Supplemental Methods for details). To simulate perturbations and apply system-wide disruptions to gene expression patterns, we randomly selected genes as perturbation targets and subsequently sampled scalar multipliers from a *Γ*(*2,2*) distribution to alter average expression values for each disrupted gene. To generate distinct cell types, we randomly selected a separate set of genes to act as markers for each population. Selected marker genes had their average expression shifted by a scalar multiplier following a *Γ*(*3,3*) distribution to create population substructure.

### ICAT analysis in real datasets

To assess ICAT’s ability to correctly cluster cells in real perturbation experiments, we made use of three publicly available datasets (Tian *et al*., 2019; Kagohara *et al*., 2020; Kang *et al*., 2017). While no true benchmark datasets currently exist for defining cell states in perturbation experiments, we selected datasets that either contain genotyped cell lines, or are from well-studied cell types with confidently labeled cells. Because NCFS feature-weighting is a computationally intensive task, and since the Kagohara and Kang datasets each contain > 20,000 cells, it was necessary to subselect cells for feature weighting (Fig. S3). To this end, ICAT selects a representative sample of cells via submodular optimization using the *apricot* (Schreiber *et al*., 2020) Python package. Submodular optimization via *apricot* was able to effectively represent the data space using only a small number of cells (Fig. S4).

We further validate ICAT’s predictions *in vivo* using a newly generated sea urchin scRNA-seq dataset consisting of one control and two experimental conditions. Treatment-unique cell states were first identified using ICAT, then empirically assessed using quantitative single molecule fluorescence in situ hybridization (smFISH) to evaluate both the cell state-defining expression patterns and the predicted changes in relative cell state abundance between treatments.

### Benchmarking ICAT’s performance

To benchmark ICAT against other general workflows for identifying cell states in multi-condition scRNA-seq experiments, we compared four general approaches: (1) ICAT, (2) integrating datasets across treatments followed by clustering cells via Louvain clustering, (3) integrating datasets across treatments followed by clustering cells via ICAT, and (4) naive clustering where we tested the “No Integration” scenario in which cells were clustered using Louvain clustering without accounting for treatment status (Fig. S4). Each workflow focuses on identifying cell states, making them intrinsically comparable despite algorithm differences.

In our evaluation we tested three state-of-the-art integration methods: Seurat (Stuart and Satija, 2019; Satija *et al*., 2015), Scanorama (Hie *et al*., 2019), and Harmony (Korsunsky et al. 2019). These methods were chosen due to their ubiquity in single cell analysis, as well as their top performance in rigorous benchmarks comparisons (Luecken *et al*., 2022). Accordingly, we also evaluated clustering performance for three extended workflows: ICAT_Seurat,_ ICAT_Scan_, and ICAT_Harm_ whereby data from cells was first integrated using the respective integration algorithm prior to clustering with ICAT. While ICAT is relatively robust to parameter choice (Fig. S5), we used standardized nearest neighbor and resolution parameters for each method during clustering for appropriate comparisons (Supplemental Methods).

When evaluating cell state labels produced in simulated experiments, we considered three criteria reflecting reliable clustering between control and treated cells: 1. Global label conservation: to ensure cluster labels accurately reflect the underlying biology, we used the Adjusted Rand Index (ARI) to measure the extent of mapping between cluster labels and known cell states (Hubert and Arabie, 1985). 2. Treatment mixing: the local homogeneity of shared cell types without undo isolation by treatment status was assessed with the Local Inverse Simpson’s Index (Korsunsky *et al*., 2019) (LISI); 3. Unique label separation: the accurate detection of populations of interest was measured using population-specific F1 measures (Luecken *et al*., 2022). Such metrics were used to measure the ability to accurately detect asymmetric populations that exist only in single conditions (F1-Unique), the ability to discriminate stimulated cell states from non-stimulated analogues (F1-Stim), and the identification of rare cell types (F1-Rare). ARI, LISI, and all F1 metrics were standardized between 0 and 1, where 1 indicated stronger performance.

Because cell state is resolution dependent and likely imperfect in real datasets (Luecken and Theis, 2019; Lähnemann et al., 2020), when evaluating performance using real data, we also calculated the label-free Davies-Bouldin (DB) metric to assess the density of clusters produced by each method (Davies and Bouldin, 1979). The DB metric measures similarity between a cluster and the next most similar cluster. To facilitate comparison, each score was scaled between 0 and 1 such that 1 represents better clustering while 0 represents worse clustering.

## Results

### ICAT accurately clusters cells in perturbation experiments regardless of compositional asymmetries

To validate the general ICAT workflow (Fig. 1), we simulated data to compare ICAT to naively clustering cells using Louvain community detection without accounting for treatment status (Fig. 2A). To initially validate the general approach of ICAT, we first performed a set of simple experiments consisting of equal size populations to compare ICAT to naive Louvain clustering (Fig. 2A, top row). Subsequently, we tested four different simulated experiments with each condition consisting of a different number of shared and unique populations between control and treated cells (Table S1). ICAT exhibited near perfect clustering with ARI scores near 1 across five different simulated experiments (Fig. 2B). Without accounting for the systematic noise introduced by perturbations, naive Louvain clustering erroneously separates cell states by treatment status (Fig. 2A) and only produced accurate cluster labels when no shared cell states between control and treated cells existed (Fig. 2B). These results not only demonstrate the necessity of accounting for treatment status while identifying cell states in perturbation experiments, but also provide validation for ICAT’s novel approach.

### ICAT is robust to perturbation severity and better resolves cell states at lower resolutions than leading integration workflows

To further challenge ICAT’s ability to robustly detect cell states in more complex scenarios, we simulated five populations of cells: three that are present in both control and treated cells, with one of these exhibiting elevated marker gene expression upon treatment and referred to as “stimulated” (population C1 and P(C1)+), and two treatment-only populations (populations P4 and P5) (Fig. 3A-B). These populations were simulated under varying conditions to assess ICAT’s robustness to varying conditions, such as perturbation severity, degree of cell state separation, and different population abundances.

We first assessed ICAT’s robustness against increasing perturbation severity, testing five conditions with varying proportions of perturbed genes, ranging from 1% to 25% (Fig. 3C, Table S2). Next, we assessed performance with changing resolutions between cell states by simulating 10 different experimental conditions with a varying number of average marker genes per cell type. Mean marker gene numbers ranged from 10-105 per population (0.67-7.0% of total genes) (Fig. 3D, Table S3). In both cases, each experiment was simulated 15 times, for a total of 90 independent datasets in the first case (Fig. 3C) and 150 in the second case (Fig. 3D). Finally, to test performance as cell type proportions deviate from equality, we simulated three different experimental conditions with varying distributions of each cell type. Experiments ranged from the most similar, with all cell types having 100 cells per treatment, to the most disparate where the minimum and maximum population size per treatment were 50 and 150, respectively (Fig. 3E, Table S4). To isolate the effect of differential cell abundance, proportion simulations did not include an activated cell type, and instead were composed of three shared populations and two treatment-unique populations (Fig. 3E).

When we tested perturbation severity or variable numbers of marker genes (Fig. 3C-D), ICAT outperformed both Scanorama and Seurat across all metrics while exhibiting highly stable performance across both increasing perturbation intensities and at varying cell-state resolutions (Fig. 3F). ICAT provides near-perfect treatment mixing, with LISI scores nearing 1 for all signal levels. In contrast, Scanorama and Seurat only approach 0.5 LISI scores at the highest signal levels. Harmony generally produces higher LISI scores than both Scanorama and Seurat, but displays significant performance loss as perturbation severity increases, while failing to match ICAT performance across cells state resolutions, indicating that ICAT exhibits a substantially improved ability to correctly and robustly mix cell states across treatments, even at low signal levels at which the standard integration algorithms fail (Fig. 3F).

Scanorama offered comparable performance to ICAT in global label conservation (ARI) and in the ability to detect asymmetrical populations (F1-Unique) (Fig. 3F), while ICAT was uniformly better at accurately distinguishing the stimulated population from its non-stimulated analogue (Fig. 3F, F1-Stim). Seurat underperformed both Scanorama and ICAT in label conservation and asymmetrical population detection (ARI, F1-Unique), and offered comparable treatment-mixing performance to Scanorama (LISI), indicating that Seurat does not accurately separate known populations relative to Scanorama or ICAT (Fig. 3F). Harmony displayed inconsistent performance, showcasing dependence on both perturbation severity and marker gene abundance to correctly identify asymmetrical populations (F1-Stim, F1-Unique).

For each of the perturbation severity tests and with low marker gene numbers, Seurat and Harmony were surprisingly outperformed by the no-integration scenario for ARI (Fig. 3F) which can be explained by both methods’ poor identification of asymmetrical populations (F1-Unique and F1-Stim, respectively) reflecting a general under-clustering of cells (Fig. S6). Scanorama exhibited a similar drop in ARI performance at low marker numbers, but consistently outperformed no integration (Fig. 3F). These results show ICAT more accurately and robustly resolves cell states across experimental conditions than either Seurat or Scanorama integration workflows.

### ICAT most accurately detects rare populations

ICAT was unaffected by differences in cell state proportions as assessed by global and rare population label conservation via ARI and the F1-Rare metric, respectively, while each integration approach exhibited decreased performance as the disparity in proportions increased (Fig. 3G, left). The performance of ICAT was robust and stronger than both Scanorama, Seurat, and Harmony in each measure.

### ICAT improves and stabilizes the performance of integration workflows

Clustering cells in integrated datasets with ICAT, in lieu of traditional Louvain community detection, produced overall higher quality clusters compared to standard integration workflows. Compared to the typical Seurat and Harmony workflows, ICAT_Seurat_ and ICAT_Harm_ led to significant improvements in all three simulated experiments, leading to greater robustness to perturbation (Fig. 3F1, left), increased signal sensitivity (Fig. 3F2, left), and better isolation of rare cells (Fig. 3G, right). When compared to Scanorama, ICAT_Scan_ better isolated rare cells (Fig. 3G, right), and improved performance in some metrics in perturbation and signal experiments. Gains were more modest compared to ICAT_Seurat_ and ICAT_Harm_, primarily since Scanorama with Louvain clustering already exhibits strong performance (Fig. 3F1-2, left). In particular, ICAT_Scan_ substantially improved treatment-mixing (LISI) across conditions compared to Scanorama with Louvain (Fig. 3F1, right). Interestingly, at both signal extremes, stand-alone ICAT scored higher in the F1-Stim metric than either combined or integration method, whether combined with Louvain or ICAT for clustering (Fig. 3F2), indicating that ICAT preserves discriminatory signals better than either integration algorithm.

In summary, the results from simulated data demonstrate that ICAT offers robust and accurate cell state identification that exceeds three current state-of-the-art integration algorithms, across a range of simulated conditions including severe perturbations, low marker gene numbers, and variations in relative cell abundance. Further analysis shows that identifying cell states with ICAT following integration produces higher quality clusters compared to current integration workflows alone. Thus, the novelty of using learned feature weights combined with control-guided clustering offers a significant performance improvement over state-of-the-art integration approaches in accurately defining the cellular compositions of single-cell datasets; moreover, the results show that, unlike integration approaches, ICAT’s performance is robust across a range of conditions.

### ICAT produces higher quality clusters in real datasets compared to integration methods

To ensure ICAT produced clusters that accurately reflect known biological cell states, we evaluated ICAT in three separate real scRNA-seq datasets. We first used the CellMix dataset generated by Tian et al. (Tian *et al*., 2019). The dataset that was produced by creating various mixtures of nine cells from combinations of H2228, HCC827, and H1975 adenocarcinoma cell lines, sequencing the cell mixtures, then downsampling the resulting counts to produce pseudo-cells that are in-line with true single-cell data. While the Tian dataset was developed to test integration across batch effects, we reasoned that variance introduced by batch effects offers an imperfect substitute for a perturbation. To better mimic distinct phenotypes, we selected only four cell “types”, consisting of pure H2228, HCC827, and H1975 cells, along with a single mixture with equal proportions of each cell line, denoted “Mixed” (Fig. S7, Table S5). The second dataset from Kagohara et al. (Kagohara *et al*., 2020) offers a simple perturbation experiment in which three genotyped cell lines of head and neck squamous carcinoma cells were treated with either PBS (control) or the chemotherapeutic cetuximab (Table S5). Finally, we assessed performance in a peripheral blood mononuclear cell dataset collected by Kang et al. (Kang *et al*., 2017) with eight cell types in control and IFN-β-stimulated conditions (Table S5). Both the Tian and Kagohara datasets were chosen for having gold-standard cell labels via genotyping, while the Kang dataset was selected to include a more complex, but well-studied, system with silver-standard cell labels identified via known marker genes. Prior to analysis, each dataset was preprocessed by normalizing cell counts and selecting highly variable genes using *Scanpy*. To ensure each tested algorithm analyzed the same input data, we provided the same *Scanpy* preprocessed data to each method.

In the Tian dataset, ICAT outperformed Seurat and Scanorama in LISI and DB metrics and was comparable to both in ARI, where Harmony and Seurat scored highest (Fig. 4A-B, Table S6). ICAT outperformed all three integration methods in the Kagohara and Kang datasets (Fig. 4A-B). Aside from ARI in the Kagohara dataset, in which Scanorama and ICAT scored equally well (Fig. 4B), ICAT produced the highest scores for ARI, LISI, and DB metrics (Fig. 4B). Overall, ICAT generated higher quality clusters in real datasets compared to existing integration algorithms (Fig. 4A-B, Table S6).

**Figure 4:**
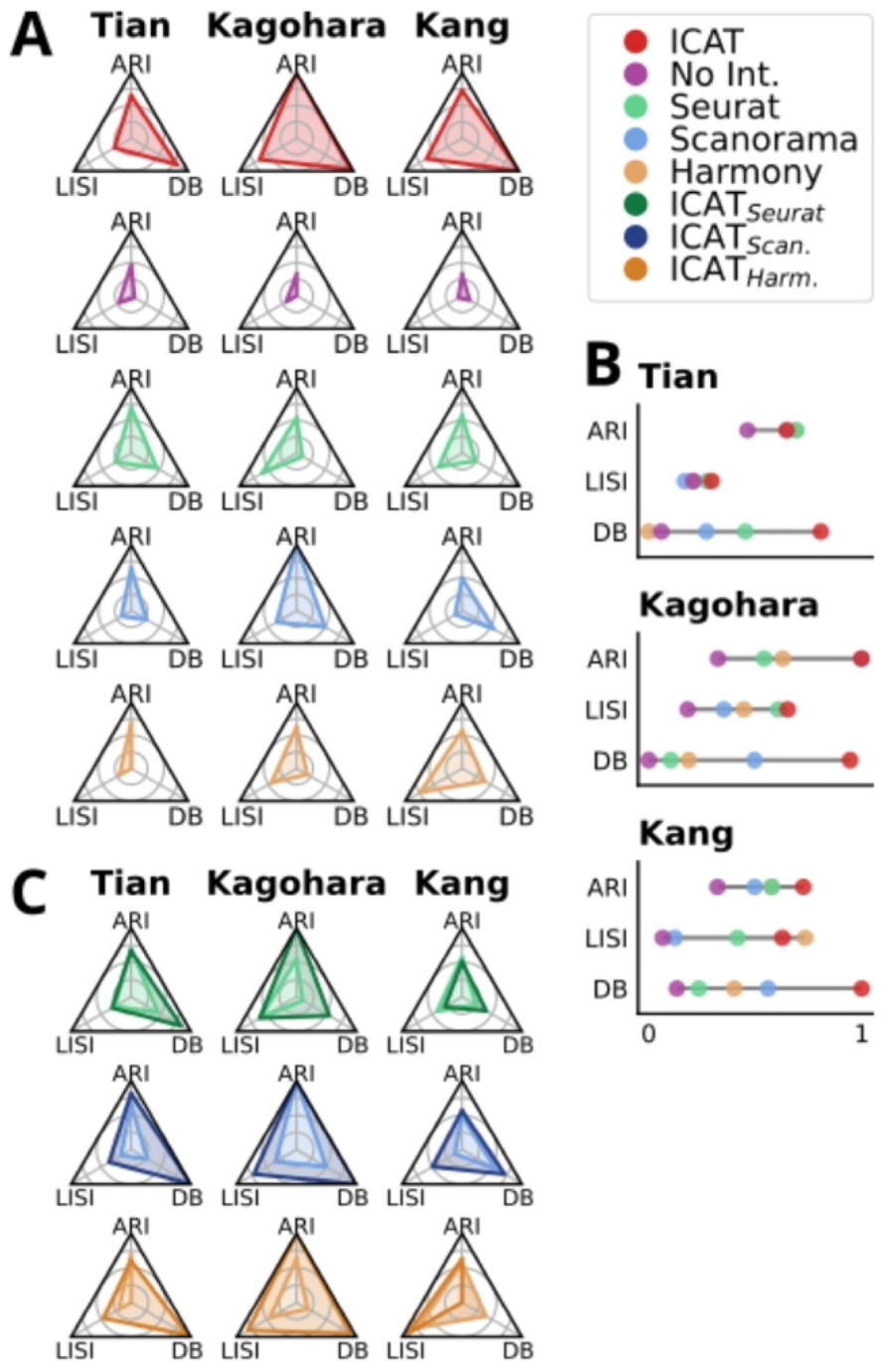
ICAT outperforms current integration methods in identifying cell states across treatments in real datasets. scRNA-seq data from Tian, Kagohara, and Kang studies is compared. **A**. Spider plots compare the performance for each algorithm within each dataset for ARI, LISI, and DB quality metrics. **B**. Lollipop plots highlight the differences in the metrics for each method across the same datasets. **C**. Spider plots comparing Seurat to ICAT_Seurat,_ Scanorama to ICAT_Scan_, and Harmony to ICAT_Harm_, respectively. All metrics are scaled from 0 to 1, where 1 is best (closest to the apex of each corner). Abbreviations: DB Davies-Bouldin metric, and otherwise as in Fig. 2. ICAT performance for the Tian dataset was evaluated using the ICAT_C+T_ implementation, while Kagohara and Kang datasets were evaluated using the ICAT_C_ implementation.

Compared to Scanorama, ICAT_Scan_ led to greater ARI, LISI and DB scores across the three analyzed datasets (Fig. 4C, Table S6), aside from ARI in the Kagohara dataset where Scanorama and ICAT_Scan_ scored equally well (Fig. 4C). For Seurat and Harmony, ICAT_Seurat_ and ICAT_Harm_ resulted in improvements that exhibit dataset and metric dependence. ICAT_Seurat_ and ICAT_Harm_ more readily identified cell states in the Kagohara dataset, shown by the dramatic increase in ARI scores, and clusters were tighter and better separated as shown by large increases in DB scores (Fig. 4C). In contrast, ARI scores were maintained or marginally reduced compared to traditional Seurat + Louvain or Harmony + Louvain workflows for the Tian and Kang datasets. The results demonstrate that ICAT offers a marked improvement in cell state identification in multi-condition scRNA-seq experiments compared to the current integration workflows Seurat, Harmony, and Scanorama. Thus, the performance of ICAT with real data is substantially improved compared to conventional integration + clustering workflows, in agreement with ICAT’s performance in simulated data.

### ICAT alone accurately predicts subpopulation response to perturbation *in vivo*

To empirically test ICAT predictions on cell state compositions following perturbation, we generated scRNA-seq data using the SMART-Seq2 protocol (Picelli *et al*., 2014). Specifically, we isolated and sequenced the skeletogenic lineage in sea urchins known as primary mesenchyme cells (PMCs) at 18 hours post-fertilization to compare cells from control embryos and embryos subjected to chemical treatments that perturb skeletal patterning, specifically, MK-886 (MK) and chlorate (Pidgeon *et al*., 2007; Simionato *et al*., 2021; Piacentino *et al*., 2016). PMCs are of interest because this population of cells receives patterning cues during development that cause their diversification into subsets; since MK and chlorate each perturb distinct patterning cues, we reasoned that these differences would be reflected in PMC subpopulation composition (Piacentino *et al*., 2016; Lyons *et al*., 2014; Sun and Ettensohn, 2014; Piacentino *et al*., 2015). After quality control and cell filtering, we retained expression data for 435 cells (147 control, 134 MK886, and 154 chlorate-treated cells).

ICAT clustered the combined PMC expression profiles into five distinct cell states (Fig. 5A1). The impacts of MK exposure were more dramatic than those of chlorate (Fig. 5A2); we therefore focused on the effects of MK. Two clusters showed stark compositional differences between control and MK-treated cells, in that cluster 3 lacks MK cells compared to controls, while clusters 2 and 4 show significant enrichment for MK cells (G-test fdr < 0.05, post hoc Fisher’s Exact test fdr < 0.05; Fig. 5A2, Table S7-8). Since MK-treated cells dominated cluster 4 such that it appears to be an asymmetric population, we more closely examined clusters 3 and 4. Investigating highly predictive genes identified by ICAT found that clusters 3 and 4 were distinguished by inverse expression of genes *pks2* and *sm50* (Fig. 5A3-4), such that cluster 3 displayed a *sm50+/pks2-*phenotype, while cluster 4 exhibited a reciprocal *sm50-/pks2+* pattern.

**Figure 5:**
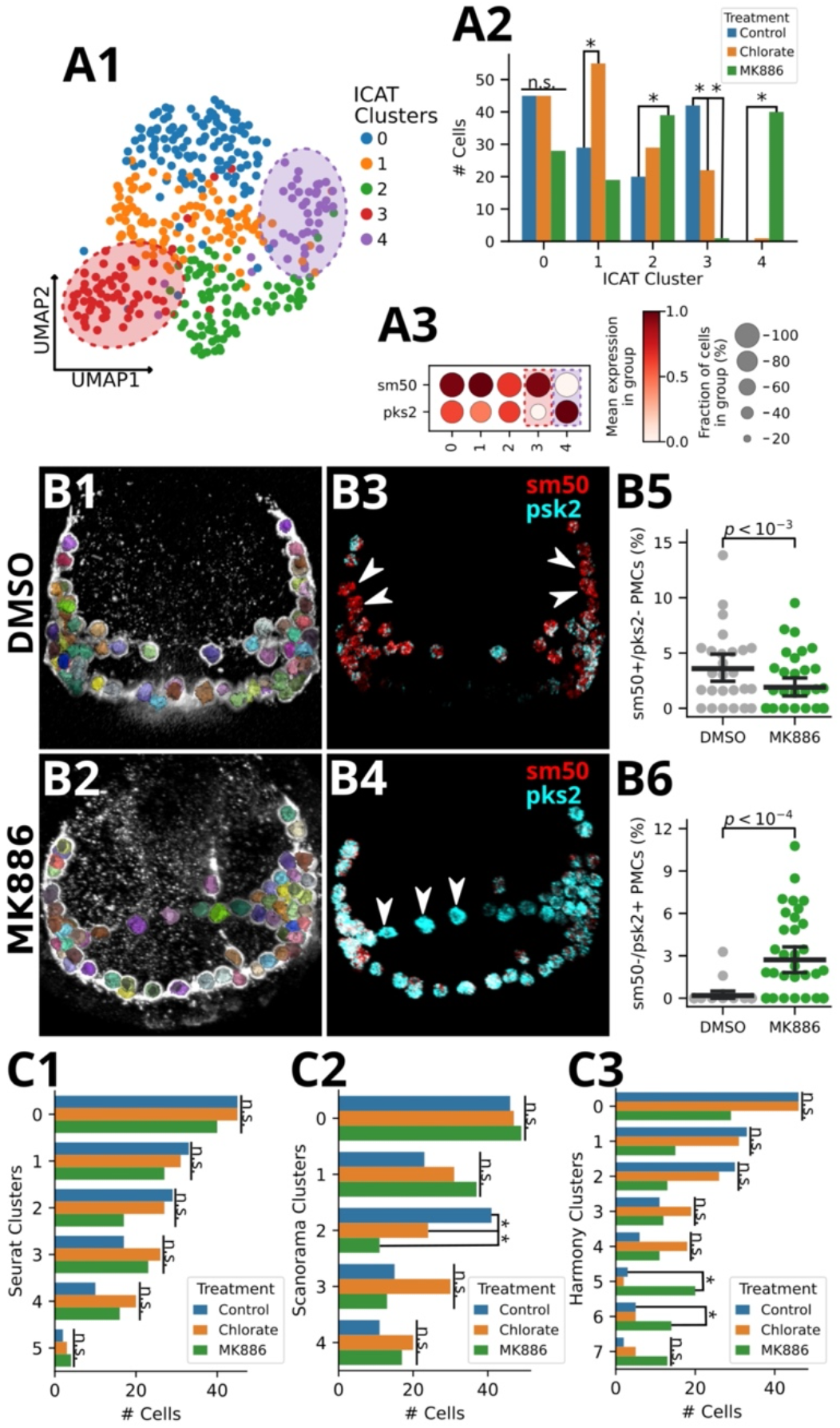
ICAT most accurately defines subpopulation response to perturbation. SMART-seq2 was performed on cells isolated from controls or from embryos treated with the perturbants chlorate or MK-886. **A**. ICAT predicts five clusters from the combined data (A1), with treatment-dependent subpopulation compositions (G-test likelihood test, fdr < 0.05; ad-hoc pairwise Fisher’s Exact test, fdr < 0.05) (A2). ICAT-calculated gene weights plotted by rank show *sm50* and *pks2* as highly informative for cell-state membership, with 37 total genes being considered informative (weight > 1) (A3). MK-induced differences are defined by reciprocal expression patterns of ICAT-identified genes, *sm50* and *pks2*, in subpopulations 3 (*sm50+/pks2-*) and 4 (*sm50-/pks2+*) (A4). **B**. ICAT predictions for control (B1,3,5) and MK-866 (B2,4,6) are validated by HCR FISH analysis. Automated detection and segmentation of PMCs *in vivo* from 3-D image data is shown as various colors; each color represents an individual cell (1-2). The expression of *sm50* (red) and *pks2* (cyan) transcripts are shown in the same embryos; arrowheads indicate PMCs that express only *sm50* or only *pks2* (3-4). MK-866 treated embryos exhibit statistically significantly fewer *sm50+/pks2-*PMCs (B5) and more *sm50-/pks2+* PMCs (B6) than controls, consistent with the predictions from ICAT (A). Dot plot error bars denote 95% confidence intervals of the mean with each dot representing an individual embryo (n_DMSO_ = 26, n_MK886_ = 37; B5 (Binomial GLM, *µ*_DMSO_= 3.59%, *µ*_MK886_= 1.91%; β_MK886_= -0.71, p < 10^−3^; B6 Binomial GLM, *µ*_DMSO_= 0.19%, *µ*_MK886_= 2.71%; β_MK886_= 2.60, p < 10^−4^). **C**. Seurat (C1) fails to find any treatment effect on PMC subpopulation composition, while Scanorama (C2) and Harmony (C3) predict one and two disrupted populations, respectively (G-test likelihood test, fdr < 0.05; ad-hoc pairwise Fisher’s Exact test, fdr < 0.05).

To empirically test ICAT’s predicted enrichment of the *sm50-/pks2+* cell state and depletion of *sm50+/pks2* cells in MK886-treated PMCs, we performed single molecule fluorescence *in situ* hybridization (smFISH) to quantify expression levels for both genes in DMSO (control) and MK-treated embryos (Choi *et al*., 2018) (Fig. 5B, Fig. S8). To accurately map FISH signals to PMCs, embryos were stained for PMCs using the PMC-specific 6a9 antibody. Using Ilastik (Berg *et al*., 2019), we trained a random forest classifier (out-of-bag error = 0.14) to predict and segment individual PMCs in each confocal image stack (Fig. 5B1-2, Fig. S9). Following methods for relative FISH quantification from the original paper (Choi *et al*., 2018), we calculated cell-level expression values for each PMC by first restricting fluorescence signal to PMC-labeled voxels, then calculated the average relative intensity for each gene in every PMC (Fig. 5B3-4, Fig. S8). Across embryos, we labeled individual PMCs as *sm50+* or *sm50-*if their standardized *sm50* expression values were above the 75th or below the 25th percentiles of all PMCs, respectively (Fig. S10). The procedure was repeated for *pks2* to determine *pks2+* and *pks2-*PMCs. We found that MK-treated embryos displayed both a significant loss of the *sm50+/pks2-*phenotype, as well as a significant enrichment of the *sm50-/pks2+* cell state relative to controls (adjusted p < 0.05; Fig. 5B5-6, Table S9-10). These results empirically validate the ability of ICAT to correctly identify affected cell states in scRNA-seq datasets from perturbed samples.

Integrating the PMC scRNA seq data using Seurat, Scanorama, or Harmony failed to identify both *sm50/pks2*-defined cell states affected by MK-886 treatment (Fig. 5C, Fig. S11). In fact, Seurat failed to identify any dependence of cell state composition on treatment status (G-test fdr > 0.05; Fig. 5C1, Fig. S11A1, B1, Table S11). Scanorama, on the other hand, correctly identified the depletion of *sm50+/pks2-*PMCs in MK embryos (Fig. S11B2-3, Table S12-13; G-test fdr < 0.05, post hoc Fisher’s Exact test fdr < 0.05), but importantly, failed to identify *sm50-/pks2+* enrichment in MK embryos (Fig. S11B2-3, Table S12-13; G-test fdr > 0.05). Conversely, Harmony correctly identified the enrichment of sm50-/pks2+ PMCs in MK embryos but lacked the sensitivity to identify the depletion of sm50+/pks2-cell states (Fig S11B3, Table S14-15; G-test fdr < 0.05, post hoc Fisher’s Exact test fdr < 0.05). ICAT is the only method among these four to correctly identify the cell-state specific *sm50* and *pks2* responses to MK exposure, which accounts for most of the differences in cell groupings between the methods (Jaccard similarity, Fig. S11C). Of the compared methods, Seurat failed to produce even qualitatively correct results since it did not identify any treatment-specific effects on PMCs (Fig. S11A2), contradicting the known disruptions to PMCs in response to disruption of the ALOX signaling pathways (Piacentino *et al*., 2016).

### ICAT-selected marker genes reflect underlying biology

By leveraging weighted expression of learned cell state-defining genes, ICAT is able to more correctly label cells in mixed-condition scRNA-seq experiments compared to traditional integration workflows. A latent assumption in this process is the relative stability of marker genes in non-treatment affected cell states between conditions. To test this assumption in real data, we performed differential expression (DE) analysis between treatments using MAST (Finak et al., 2015) (Fig. S12). In the Tian and Kagohara datasets, none of the highly weighted genes were found to be DE (log_2_ fc > 1, fdr < 0.05, Fig. S12), while only a single gene in the Kang dataset was considered DE when a looser log_2_ fc threshold was applied (log_2_ fc > log_2_(1.5), fdr < 0.05, Fig. S12). These results are consistent with the assumption of marker gene stability across perturbations. However, importantly, four marker genes were considered DE in the PMC dataset (log_2_ fc >1, fdr < 0.05, Fig. S12); these genes contributed to ICAT’s ability to sensitively identify the verified sm50/pks2 perturbed cell states. These outcomes underline the robustness of ICAT when challenged by marker genes whose expression is variable. Together, these results demonstrate that ICAT’s novel combination of learned feature weighting combined with control-guided clustering significantly improves the computational capability to accurately detect and describe cellular responses to perturbation in scRNA-seq data compared to conventional integration workflows.

## Discussion

One of the key promises of scRNA-seq technology is to more accurately define complex biological systems by offering a more finely resolved perspective on transcriptional landscapes than was previously possible. However, due to technical and biological noise, taking advantage of such a powerful tool in diverse biological conditions has proven to be challenging. We therefore developed the ICAT algorithm to improve the accuracy of cell state identification from scRNA-seq data that includes perturbation treatments. By combining sparse feature weighting with semi-supervised Louvain community detection, ICAT not only correctly identifies shared and disparate cell states, but also produces interpretable results that aid in understanding analytical outcomes and selecting the most suitable downstream analyses.

When comparing the performance of ICAT to state-of-the-art integration algorithms in simulated datasets, we found that ICAT performs more robustly in response to perturbation severity, readily identifies cell states at lower signal levels, better isolates asymmetrical and activated cell states, and more readily identifies rare cell populations when compared to Seurat, Scanorama, and Harmony. ICAT also outperforms the integration workflows in three benchmark datasets, further solidifying ICAT’s ability to better identify cell states across biological conditions compared to either integration method alone. By testing ICAT on datasets with > 20,000 cells (Kang *et al*., 2017; Kagohara *et al*., 2020), we demonstrated ICAT’s ability to analyze the large datasets, necessary for any practical scRNA-seq tool. Depending on the experimental system, researchers may still elect to use integration algorithms. Excitingly, the choice between ICAT and integration methods is not mutually exclusive; we show that following Seurat, Scanorama, or Harmony with ICAT produces higher quality clusters compared to either of these integration workflows alone.

By experimentally validating two uniquely predicted cell states in a developmental model using single molecule FISH, we empirically demonstrated that the capability of ICAT to accurately identify perturbed cell states surpasses current state-of-the-art integration workflows. ICAT was the only method among the four tested that correctly identified these distinct states and their perturbation responses, whereas integration workflows lacked the sensitivity to properly isolate both sm50/pks2 defined subpopulations, highlighting ICAT as a substantial advancement in identifying and describing cell state behaviors across biological conditions and perturbations.

With the growing ubiquity of scRNA-seq experiments and the development of ever more sophisticated experimental designs, we anticipate that ICAT will play an important role in highly resolving cell state-specific responses to treatments and differing biological conditions. In the future, the ICAT framework can easily be extended to operate on other single-cell modalities, such as scATAC-seq, and will offer a potential route to intelligently cluster multi-modal data from heterogeneous conditions.

## Supporting information

Hawkins et al. Supplemental Material

## Declarations

### Availability of data and materials

*ICAT, ncfs*, and *sslouvain* Python packages are freely available on pip and github (github.com/BradhamLab). Scripts to perform simulated data generation and benchmark comparisons are available as a SnakeMake workflow at (github.com/BradhamLab/ICAT_manuscript). Archived data files for both simulated and real data are also available at the same repository. The pipeline to align and quantify SMART-seq2 reads is freely available at (github.com/BradhamLab/scPipe), and the raw sequencing data is available at GEO accession GSE164240. Likewise, all scripts to segment PMCs and quantify HCR FISH signal are available at (github.com/BradhamLab/hcr_quantify) and the image data is available upon request.

## Acknowledgements

We thank Prof. Masanao Yajima (Dept. of Mathematics and Statistics, Boston University) for helpful discussions. We thank Prof. Charles Ettensohn (Carnegie Mellon University, PA, USA) for the PMC antibody. We thank Jose Ordovas-Montanes and Sanjay Prakadan of the Shalek lab (MIT) for help with scRNA-seq library preparation. We thank Dr. Todd Blute for his advice and help with FACS and microscopy imaging.

## Author Contributions

The study was conceived by CAB and DYH. The computational work was performed by DYH, with advice from WEJ. JH and DTZ collected the urchin scRNA-seq data. NRS, CFT, KRM, ATL, and AED performed the imaging experiments. DYH, KRM, and AG trained the PMC prediction model. The manuscript was written by DYH, CAB, DTZ and NRS, and edited by all coauthors.

