## Supplementary material for "ICAT: A Novel Algorithm to Robustly Identify Cell States Following Perturbations in Single Cell Transcriptomes": Hawkins et al. Supplemental Material

**Hawkins et al. 2023 Supplemental Materials**

**A. Supplemental Methods**

**B. Supplemental Tables 1-16**

**C. Supplemental Figures 1-12**

### Supplemental Methods

#### Initial clustering

Prior to feature weighting, ICAT identifies initial clusters using Louvain community detection via the Scanpy Python package (Blondel *et al.*, 2008; Wolf *et al.*, 2018). The neighbor graph is built using default parameters of *scanpy.pp.neighbors* with the exception of *n_neighbors* and *resolution* parameters, which are optimized for each dataset. Following Scanpy best practices, cell-cell distances are measured using principal components calculated using the *scanpy.pp.pca* function, again with default parameters. Once the neighbor graph is constructed, ICAT performs Louvain community detection using the *scanpy.tl.louvain* function.

#### Neighborhood Component Feature Selection

Neighborhood Component Feature Selection (NCFS) was implemented in Python following the methods outlined in the original publication (Yang *et al.*, 2012). The implementation takes advantage of the just-in-time compiler library, NUMBA, to produce highly performant code (Lam *et al.*, 2015). NCFS optimizes feature weights to maximize the score function:

$$\xi(\mathbf{w}) = \sum_i \sum_j y_{ij} p_{ij} - \lambda \sum_{l=1}^{d} w_l^2$$

where $\mathbf{w}$ is the weight vector; $y_{ij} = 1$ if samples $i$ and $j$ belong to the same class, and is 0 otherwise; $p_{ij}$ is the probability of selecting $j$ as a reference to $i$ in a kNN classifier; and $\lambda > 0$ is a regularization parameter. We alter the score function by including a class weight term, $c_i$, for each sample to account for class imbalance:

$$\xi(\mathbf{w}) = \sum_i c_i \sum_j y_{ij} p_{ij} - \lambda \sum_{l=1}^{d} w_l^2$$

Sample weights are calculated using methods outlined in the scikit-learn Python package (Pedregosa *et al.*, 2011; King and Zeng, 2001). Gradient ascent is then performed using the line method described in the original publication.

**Semi-supervised Louvain community detection**

To identify partitions in a graph, traditional Louvain community detection initializes each node into singleton clusters. Nodes are then merged into other communities if doing so would optimize a specified quality score (Blondel *et al.*, 2008). To perform semi-supervised clustering, ICAT initializes communities with previously identified control cluster labels. Nodes representing perturbed cells are placed into their own singleton clusters. During optimization, initialized control labels are set immutable, while labels for perturbed cells are free to change. By keeping control labels immutable, we simultaneously allow perturbed cells to merge into control communities — to identify mixed cell states — while also allowing isolated nodes to form treatment-specific communities. By default, ICAT optimizes graph partitions using the Reichardt-Burnholdt metric (Reichardt and Bornholdt, 2006). Semi-supervised Louvain was implemented by extending the popular louvain-igraph package (github.com/vtraag/louvain-igraph) to allow for immutable nodes.

**Subsampling by submodular optimization**

Due to the computational intensity of NCFS feature weighting, for large datasets it is beneficial to learn feature weights on a subset of the overall cells. To select cells that correctly capture the dataset variation, ICAT employs submodular optimization via the *Apricot* Python package (Schreiber *et al.*, 2020). By default, ICAT uses the *facilityLocation* model in *Apricot* to successively select cells that represent underrepresented regions of the data. The number of cells to select is chosen by the user; this was set to 1,500 for both Kagohara and Kang datasets.

**Simulating scRNA-seq perturbation experiment data**

Simulated data was generated using a zero-inflated negative binomial distribution to model gene counts. Our approach follows methods outlined by Büttner and colleagues (Büttner *et al.*, 2019), used to simulate batch effects in scRNA-seq data, adopted to provide for multiple cell populations as well as different cell identity resolutions and perturbation severity.

For each dataset, we simulated 1,500 genes. Letting $G = \{g_1, ..., g_{1,500}\}$ be the set of all genes, to model $M$ distinct cell identities, we simulate $K$ marker genes such that $G_k = \{g^1, ..., g^K\} \subset G$ and $K \sim B(n = 1500, p = p_{marker})$, where $p_{marker}$ is set for each simulation. For simplicity, marker genes were made to be mutually exclusive, such that a gene could not be selected as a marker for multiple populations. Count matrices were constructed as follows:

Denote each cell identity $y^k$, where $Y = \{y^1, ..., y^M\}$ is the set of all identity labels. Assuming $N$ cells, let $y_i$ be the cell identity label for cell $i$ such that $y_i \in Y$ and $G_{y_i}$ is the set of marker genes associated with the given identity. For control cells, to generate the count matrix $X$, we simulate the cell-gene expression values $X_{ij} \sim NB(\mu_{ij}, r = r_j | p_{ij})$, where $r_j \sim U(1, 4)$ is the gene-wide dispersion factor and $p_{ij}$ is the cell-gene specific dropout probability. Prior to marker gene application, gene-wide expected values are simulated using a Beta distribution, where $\mu_j \sim Beta(a = 2, b = 5) \cdot c$, where $c = 100$ is a scalar constant. For marker genes, this scalar $c$ is modified to a cell-identity specific value where $c_{y_i} \sim \Gamma(a = 3, b = 3) \cdot 100$. Thus, cell-gene expected values are modeled:

$$\mu_{i,j} = \begin{cases} \beta(a = 2, b = 5) \cdot 100, & g_j \notin G_l \\ \beta(a = 2, b = 5) \cdot c_{y_i}, & g_j \in G_l \end{cases}$$

Letting $\mu_{y_i} = \{\mu_{y_i,1}, ..., \mu_{y_i,M}\}$ be the set expected values for all gene expression values for cells in population $y_i$, the cell-gene specific dropout probability, $p_{ij}$ is set using the sigmoid function $p_{ij} = sigm\left(-\beta_0 + \beta_{y_i j} \cdot \mu_{ij}\right)$, where $\beta_0 = -1.5$ and $\beta_{y_i j} = median(\mu_{y_i})^{-1}$.

To simulate the count matrix for perturbed cells, $X^{pert}$, we use the general framework described above, but apply a scalar shift to gene-wide expected values for perturbed genes. First, genes to perturb are randomly sampled from $G$ without replacement, such that $G^{pert} \subset G^{sim}$ is the set of all genes to perturb, and $G^{sim}$ is the set of all allowable genes to perturb. By default, $G^{sim}$ is the set of all non-marker genes such that $G^{sim} = G \setminus \bigcup_{k=1}^{k=M} G_k$. When simulating a stimulated population, we specifically included all marker genes for target population $y_{stim}$ in $G^{pert}$ where the rest of the genes were again sampled from $G^{sim}$. Assuming gene $g_j \in G^{pert}$, a perturbation scalar is sampled from a Gamma distribution such that $s_j \sim \Gamma(a = 2, b = 2)$. To simulate expression changes due to perturbation, this scalar is multiplied by cell-gene expected values. Letting $\mu_{i,j}^{pert}$ be the expected value for gene $g_j$ of cell $i$ in $X^{pert}$,

$$\mu_{i,j}^{pert} = \begin{cases} \mu_{ij}, & g_j \notin G^{pert} \\ \mu_{ij} \cdot s_j, & g_j \in G^{pert} \end{cases}$$

By plugging in $\mu_{ij}^{pert}$ into the described count simulation framework, we simulate both cell-gene dropout probabilities and the final counts. An exception is made for marker genes: in order to ensure expected population structure exists in both control and perturbed count matrices, dropout probabilities of marker genes were forced to be the same between the two conditions.

**Simulating experimental conditions**

To increase the severity of perturbation and cell identity separation, we varied $|G^{sim}|$ and $p_{marker}$, respectively. For perturbation severity experiments, $|G^{sim}|$ was set to 1%, 5%, 10%, 15%, 20%, or 25% of the total number of genes. To increase cell identity signal strength, $p_{marker}$ was chosen from the range [0.007, 0.07] with a step size of 0.07. Given a binomial distribution, the average number of marker genes ranged from 10.5 - 105 genes per population.

**Assessing performance in real datasets**

Raw count matrices of publicly available datasets (Kang *et al.*, 2017; Kagohara *et al.*, 2020; Tian *et al.*, 2019) were downloaded from GEO. All datasets were preprocessed using Scanpy, following typical workflows: first, cell counts were normalized by library size using the *scanpy.pp.normalize_total* function. Normalized counts were then log-transformed using the *scanpy.pp.log1p* function. Finally, treatment-specific highly variable genes were identified using the *scanpy.pp.highly_variable_genes* function, with "flavor=seurat" setting, and with each dataset's treatment variable set as the batch key. The union of highly variable genes was kept, and genes that were not highly variable in at least one treatment set were removed.

**Tian processing**

The Tian CellMix count matrices and metadata files were downloaded from GEO with ID GSE118767. To isolate more discrete populations, all cells not belonging to pure H2228, HCC827, and H1975 cell mixtures, or the cell mixture with equal proportions, were filtered out. Count matrices were preprocessed using *Scanpy* to remove genes expressed in less than three cells, and the dataset of origin (originally the batch confounder) was set to the treatment variable.

**Kagohara processing**

The Kagohara count matrices and metadata files were downloaded from GEO with ID GSE137524. Genes present in less than 50 cells were removed, along with cells with reads for less than 50 genes.

**Kang processing**

The Kang count matrices and metadata files were downloaded from GEO with ID GSE96583. Cells described as doublets in the cell metadata files were removed from downstream analysis. Further, genes present in less than 50 cells were removed, along with cells with reads for less than 50 genes.

**Louvain parameter selection**

In Louvain community detection, two parameters $k$, the number of neighbors, and $\gamma$, a resolution parameter, play important roles in identifying clusters. To optimize these parameters for each dataset (both simulated and real), we filtered each unintegrated dataset to control cells, and performed a grid search to find parameter pairs that optimized the Calinski-Harabarz score as implemented in scikit-learn(Caliński and Harabasz, 1974; Pedregosa *et al.*, 2011). Neighbor values were searched on the interval [5, 50] with a step size of 5. Likewise, we searched resolution values spanning [0.2, 1.25] with a step size of 0.05.

**ICAT feature weighting**

For the Kagohara and Kang datasets, feature weights were learned using only control cells (via the $ICAT_C$ implementation). Meanwhile, when analyzing all simulated datasets along with the Tian dataset, feature weights were jointly learned and subsequently collapsed across conditions using the

$ICAT_{C+T}$ implementation. Genes were considered "informative" if they had a learned weight > 1. NCFS parameters used for each dataset are listed in Table S16.

**Seurat integration**

Seurat integration was performed using version 3.1 as outlined in the vignette (https://satijalab.org/seurat/articles/integration_introduction.html). Datasets were integrated over the treatment variable specified for each dataset. Default parameters were used with the exception of specifying calculation of 50 principal components, and explicitly passing $k$ neighbors and $\gamma$ resolution parameters to standardize parameters between methods. To follow the expected Seurat workflow as best as possible, clusters were identified using the Seurat function, *FindClusters*, with the "algorithm=1" to perform Louvain community detection.

**Scanorama integration**

Scanorama integration was performed using version 1.4, and passing AnnData objects to the *run_scanorama()* function. Default parameters were used, with the exception of $k$ and $\gamma$ parameters being specified for each dataset. After Scanorama integration, clusters were identified using the *scanpy.tl.louvain* function from Scanpy

**Harmony integration**

Harmony integration was performed using version 0.1.1 in R using the *HarmonyMatrix()* function as outlined in the developer provided vignette (https://github.com/immunogenomics/harmony/blob/master/vignettes/quickstart.Rmd) (Korsunsky *et al.*, 2019). We used default parameters for the function, outside of the number of principal components, which was set to 50 to standardize between methods. Cells were then clustered using the *FindClusters* from the Seurat package, with "algorithm=1" to perform Louvain community detection, and with the $k$ neighbors and $\gamma$ resolution parameters standardized for each tested dataset.

**Quality metrics**

All metrics, with the exception of LISI, were calculated using the scikit-learn Python package. Metrics that rely on distance measures, Davies-Bouldin (DB) and LISI scores, were calculated using the euclidean distance over the first 50 principal components.

**Davies-Bouldin**

For DB, scores closer to zero represent a tighter cluster that is more separated from other clusters. To facilitate comparison between metrics for each dataset, the negative of DB scores was scaled between [0, 1] using the MinMaxScaler preprocessor from scikit-learn, where 1 now represents the best performing method.

**LISI**

LISI metrics were calculated using the "lisi" R package (Korsunsky *et al.*, 2019). The LISI score naturally falls between 0 and the number of batches/treatments in the dataset (representing total separation and perfect mixing, respectively). Because we expect mixing in only shared cell identities, PCA was first run on the complete data matrix, then cells from non-overlapping cell identities were removed, and LISI scores were then calculated using PCs from shared identities. Because LISI calculates a score for each cell, cell scores were averaged for each dataset-method comparison. To facilitate comparison to other metrics, LISI scores were scaled between 0 and 1 using the formula:

$$LISI_{scaled} = \frac{LISI - 1}{P - 1}$$

where $P$ is the number of treatment conditions in the dataset.

**Population specific F1-scores**

To assess each method's ability to detect stimulated and asymmetrical populations, we created population-specific F1-scores. For each population of interest, we identified the cluster where a plurality of cells was assigned, and denoted this cluster as $c$. For each cell within the population of interest, we created a binary vector, $\mathbf{v}$, denoting whether the cell was assigned to cluster $c$.

$$v_i = \begin{cases} 1 & y_i = c \\ 0 & y_i \neq c \end{cases}$$

where $y_i$ is the cluster label for cell $i$. Assuming perfect isolation where all cells belong to the same cluster, all entries would be 1. Creating a vector of ones: $\mathbf{u} = \vec{1}$, serves as the basis for comparison. We passed $\mathbf{v}$ and $\mathbf{u}$ to the *sklearn.metrics.f1_score* function to calculate the population-specific F1-score (Pedregosa *et al.*, 2011). In the case of F1-Unique, where multiple populations of interest exist, we calculate F1-scores for each population independently and average the scores between populations. F1-Stim was calculated based solely on the activated population.

**Differential Expression Analysis**

Differential expression analysis was performed using the *FindMarkers()* function from the Seurat package in R. For datasets with multiple treatments, differential expression analysis was run for each experimental condition, with the parameters "ident.1" set to the control identity and "ident.2" set to the experimental condition of interest. For all comparisons, "test.use" was set to "MAST" while the "logfc.threshold" and "min.pct" were set to -∞ to return all possible results for plotting. Genes were later considered differentially expressed if they had an fdr < 0.05 and a | log2(fc) | > 1, except for the Kang dataset, where the log2 fold change threshold was set to log2(1.5).

**Animals and perturbations**

Adult *Lytechinus variegatus* sea urchins were obtained from the Duke University Marine Laboratory (Beaufort, NC) or from Reeftopia (Miami, FL). Gamete harvesting and embryo culturing was performed as described (Piacentino *et al.*, 2016; Bradham and McClay, 2006). Embryos were treated with chlorate to inhibit the formation of sulfated proteoglycans as described (Piacentino *et al.*, 2016). MK-886 (Tocris) was reconstituted in DMSO, and dose–response experiments were performed to obtain the optimal working doses. The optimal MK-886 concentration was determined to be 1.5-2 µM.

**PMC preparation and FACS for scRNA-seq analysis**

Embryos were disaggregated at 18 hours post-fertilization (hpf) according to McClay and Fink (McClay and Fink, 1982), with modification as follows: Approximately 3000 embryos at 18 hpf were collected in artificial seawater (ASW) supplemented with pronase (0.66 mg/mL, Sigma) and immediately centrifuged for 1 minute at 800 x g, then resuspended in calcium-free seawater (CFSW: 450 mM sodium chloride, 9 mM potassium chloride, 48 mM magnesium sulfate, 6 mM sodium bicarbonate) containing 20 mg/mL BSA, followed by the addition of 1 volume of hyaline extraction medium (HEM) (McClay and Fink, 1982) and allowed to incubate for 4 minutes at room temperature (RT). Embryos were then centrifuged for 1 minute at 800 x g, resuspended in CFSW (McClay and Fink, 1982), triturated with decreasing diameter pipettes for 30 seconds total, then centrifuged at 1000 x g for 2 minutes to pellet the dissociated cells. Single cells were then resuspended in CFSW, centrifuged for 1 minute, then incubated in ASW containing 1 mg/mL BSA with 6a9 anti-PMC primary antibody (1:5) and anti-mouse Alexa-Fluor 488 labeled secondary antibody (1:500) for 5 minutes. Following washes, cells were suspended in CFSW containing Sytox Blue Dead Cell Stain (1:1000, ThermoFisher) to label dead cells for exclusion, then sorted using a Sony SH800 Cell Sorter. Cells were gated to exclude doublets and dead cells and collected into 96-well plates preloaded with RNAlysis buffer at one cell per well, for processing for single cell library preparation.

**scRNAseq library preparation and sequencing**

RNA quantitation and integrity were determined using a Qubit® 2.0 Fluorometer (Life Technologies) and a 2100 Bioanalyzer (Agilent Technologies). scRNAseq libraries were generated using Smart-seq2 on FACS-sorted PMCs as described (Picelli *et al.*, 2014; Trombetta *et al.*, 2014). Libraries were sequenced on an Illumina NXT platform (BU Microarray and Sequencing Resource Core Facility) using 75 bp paired end reads. We obtained a median of 1.4M reads per cell over a total of 416 cells. 144 of these cells were untreated (ASW, artificial sea water), 66 were treated with DMSO (vehicle), 206 with chlorate, and 192 with MK-886. For all reads, quality control was performed prior to alignment with fastp (Chen *et al.*, 2018) using default parameters. Roughly 97% of reads passed quality control measures with a median of ~1.3M reads per cell. Quality-controlled reads were aligned to the *L. variegatus* genome (Li *et al.*, 2020) using STAR (Dobin *et al.*, 2013), and alignments were quantified using featureCounts (Liao *et al.*, 2014). A median of 148,142 reads per cell were mapped and assigned to annotated regions within the genome. To ensure only well-sequenced cells and genes were considered, we removed cells with less than 25,000 mapped reads, then genes with reads assigned in less than five cells, using Scanpy (Wolf *et al.*, 2018). Once the final count matrix was generated, we normalized reads between cells and performed batch-correction using the SCRAN package (L. Lun *et al.*, 2016; Haghverdi *et al.*, 2018). Assuming some non-PMCs escaped FACS, we evaluated the identity of both the sorted and remainder cells from the FACS analysis for the expression of known marker genes for each major cell type (Hogan *et al.*, 2020). We then performed a final filtering step that removed sorted cells that lack expression of canonical PMC marker genes. After filtering, 119 ASW cells, 26 DMSO-treated cells, 154 chlorate-treated cells, and 134 MK-treated cells remained. For analysis, ASW and DMSO vehicle controls were combined into a single control group since the number of DMSO cells were too small to consider them as a representative sample for these comparisons and since comparisons of DMSO and ASW cells did not reveal significant expression differences. Prior to cluster analysis, genes were subsetted to highly variable genes identified in control cells using the *highly_variable_genes* function in *Scanpy*. For clustering, neighbor parameter k=15 and resolution parameter $\gamma = 1$ were used for both ICAT clustering, and subsequent Louvain Community Detection in Seurat and Scanorama workflows.

**Assessing treatment effect on cell state composition**

To assess whether cell state membership was dependent upon treatment, we performed G-tests on each cell state using the scipy.stats.*power_divergence* function in the *scipy* Python package (Sokal and Rohlf, 2013; Virtanen *et al.*, 2020). P-values were then adjusted using the *statsmodels.multitest.multipletests* function from *statsmodel* in Python, with *method=“fdr_bh”* (Seabold and Perktold, 2010). Associations were considered significant with $\alpha = 0.05$. Cell states with compositions dependent upon treatment status were further evaluated with post-hoc pairwise Fisher’s Exact test to determine which treatments affected cell state membership compared to control. Tests were performed using scipy.stats.*fisher_exact* function from scipy, with p-values again adjusted using the *multipletests* function with the same parameters.

**Single Molecule FISH: probe sets, amplifiers and buffers**

Embryos were collected at 18 hpf and fixed in 4 % paraformaldehyde. Hybridization chain reaction (HCR)-FISH was performed as described (Choi *et al.*, 2018) using probes, hairpins and buffers obtained from Molecular Instruments (Los Angeles, CA). Embryos were incubated with hairpins for 2.5 hours, then signals were captured as 3-D z-stacks using confocal microscopy.

**Immunostaining and confocal microscopy**

Immunolabeling was performed as described (Piacentino *et al.*, 2015; Bradham and McClay, 2006; Gross *et al.*, 2003). PMCs were detected using primary antibody 6a9 (1:50; from Prof. Charles Ettensohn, Carnegie Mellon University, Pittsburgh, PA, USA) and secondary goat-anti mouse antibody labeled with Alexa 546 (1:500; ThermoFisher). Hoechst 33342 (1:1000; ThermoFisher) was included as a nuclear counterstain. Confocal microscopy was performed using a Nikon-C2 confocal & Ti-E Spectral imaging system. Confocal z-stacks were projected using Napari (Nicholas Sofroniew *et al.*, 2020).

**PMC segmentation**

To isolate individual PMCs from the 3-D image stack, a random forest PMC classifier was trained using the "pixel classification" workflow from Ilastik (Berg *et al.*, 2019). The model was trained using DMSO and MK-treated embryos (n=70) to produce "PMC" and "Background" probabilities. Prior to training, images were Z-cropped to include only z-slices with PMC signal. Each slice was then contrast and intensity adjusted using the functions *exposure.adapt_hist* and *exposure.rescale_intensity* functions from *scikit-image (skimage)* (van der Walt *et al.*, 2014) library in Python. PMC probabilities were exported from Ilastik for post-processing and PMC segmentation by a custom Python script.

Briefly, exported Ilastik probabilities were first segmented using *filters.apply_hysteresis_threshold* from *skimage* with parameters "low"=0.45 and "high"=0.5. The resulting binary mask was then refined via morphological opening using *morphology.binary_opening*, again from *skimage*. The opened mask was subsequently used as seed points for watershed segmentation (*skimage.segmentation.watershed*) using the image gradients from the original data (*skimage.filters.sobel, axis=0)*. To restrict PMC labels to typical PMC sizes and to better separate dense PMC areas, individual regions with areas exceeding 600 voxels were further segmented using a stricter threshold workflow: first, PMC probabilities were smoothed using a gaussian filter (*skimage.filters.gaussian*). Second, starting with threshold $t = 0.8$, a binary mask of the smoothed probabilities was generated by finding all voxels > $t$ and performing a morphological opening. Isolated objects were then detected using *measure.label* from *skimage.* If the number of identified objects did not exceed one, the process was repeated after incrementing $t$ by 0.025 until the original region was split. The split labels were then filled back into the original larger region using watershed. To split cell segmentations with unrealistic 2-D diameters, this process was repeated for any labeled region having a diameter greater than 15 voxels as well as an area greater than 500 voxels. To correctly segment vertically stacked PMCs, any label that spanned more than seven z-slices was split after calculating the label diameter at each slice. Given that PMCs are spherical, it was reasoned local minima in diameter would indicate the end of a cell, and thus these served as break points between labels. Finally, to remove unrealistically small labels, the final segmentation was produced by removing any label that occupied less than 55 voxels.

**HCR FISH quantification and analysis**

Prior to quantification, images were filtered to remove embryos that displayed either poor stain contrast or failed to be entirely captured by the z stack. After quality control, 26 DMSO and 37 MK treated embryos were retained from the three fertilizations performed. Individual PMCs were then labeled using the previously described methodology. After label generation, *pks2* and *sm50* HCR FISH image channels were preprocessed using FISH-quant v2 (Imbert *et al.*, 2021) to scale and normalize intensities and to perform background subtraction and autofluorescence removal. Gene expression was quantified for each predicted PMC by calculating the average signal intensity as described (Choi *et al.*, 2018). Prior to analysis, PMC-specific intensities were standardized to a [0, 1] scale to allow for inter-embryo comparisons. Cells were considered to have decreased (-) or increased (+) expression for a given gene if its standardized expression fell below the 25th or above the 75th percentile, respectively. We then calculated the proportion of *sm50+/pks2-* and *sm50-/pks2+* PMCs for each embryo. Statistical analysis was performed using a general linear model to perform binomial logistic regression to assess the effect of treatment status on phenotype proportion while also accounting for the fertilization batch. The model was fit using the *statsmodels* (Seabold and Perktold, 2010) Python package using the formula *phenotypic* + *not_phenotypic* ~ *treatment* + *fertilization*, where *phenotypic* is the number of cells displaying the phenotype of interest, *not_phenotypic* is the number of cells not displaying the phenotype of interest, *treatment* is treatment status, and *fertilization* is the fertilization batch for the given embryo.

| Experiment | Perturbed Genes (%) | Markers (Mean) | Cells | # Pops. | Control Unique Pops. | Rx Unique Pops. |
|---|---|---|---|---|---|---|
| **All Same** | 25 | 15 | 400 | 2 | 0 | 0 |
| **Rx Unique** | 25 | 15 | 500 | 3 | 0 | 1 |
| **Control Unique** | 25 | 15 | 300 | 2 | 1 | 0 |
| **Both Unique** | 25 | 15 | 400 | 3 | 1 | 1 |
| **None Same** | 25 | 15 | 400 | 4 | 2 | 2 |

**Supplemental Table 1. Characteristics and parameters of treatment composition experiments.**

| Experiment | Perturbed Genes (%) | Markers (Mean) | Gini | Cells | # Pops. | Activated Pops. | Unique Pops. |
|---|---|---|---|---|---|---|---|
| **1** | 5 | 15 | 0 | 800 | 5 | 1 | 2 |
| **2** | 10 | 15 | 0 | 800 | 5 | 1 | 2 |
| **3** | 15 | 15 | 0 | 800 | 5 | 1 | 2 |
| **4** | 20 | 15 | 0 | 800 | 5 | 1 | 2 |
| **5** | 25 | 15 | 0 | 800 | 5 | 1 | 2 |

**Supplemental Table 2. Characteristics and parameters of simulated perturbation experiments.**

| Experiment | Markers (Mean) | Markers % (Mean) | Perturbed Genes (%) | Gini | Cells | # Pops. | Activated Pops. | Unique Pops. | Genes |
|---|---|---|---|---|---|---|---|---|---|
| **1** | 10 | 0.7 | 20 | 0 | 800 | 5 | 1 | 2 | 1500 |
| **2** | 21 | 1.4 | 20 | 0 | 800 | 5 | 1 | 2 | 1500 |
| **3** | 31 | 2.1 | 20 | 0 | 800 | 5 | 1 | 2 | 1500 |
| **4** | 42 | 2.8 | 20 | 0 | 800 | 5 | 1 | 2 | 1500 |
| **5** | 52 | 3.5 | 20 | 0 | 800 | 5 | 1 | 2 | 1500 |
| **6** | 63 | 4.2 | 20 | 0 | 800 | 5 | 1 | 2 | 1500 |
| **7** | 73 | 4.9 | 20 | 0 | 800 | 5 | 1 | 2 | 1500 |
| **8** | 84 | 5.6 | 20 | 0 | 800 | 5 | 1 | 2 | 1500 |
| **9** | 94 | 6.3 | 20 | 0 | 800 | 5 | 1 | 2 | 1500 |
| **10** | 105 | 7.0 | 20 | 0 | 800 | 5 | 1 | 2 | 1500 |

**Supplemental Table 3. Characteristics and parameters of simulated signal experiments.**

| Experiment | Gini | Perturbed Genes (%) | Markers (Mean) | Cells | # Pops. | Activated Pops. | Unique Pops. | Genes |
|---|---|---|---|---|---|---|---|---|
| **1** | 0 | 20 | 10 | 800 | 5 | 0 | 2 | 1500 |
| **2** | 0.12 | 20 | 10 | 800 | 5 | 0 | 2 | 1500 |
| **3** | 0.23 | 20 | 10 | 800 | 5 | 0 | 2 | 1500 |

**Supplemental Table 4. Characteristics and parameters of simulated proportion experiments.**

| Dataset | Treatments | Cell Types | Distinct Cell Types (#) | Cells (#) |
|---|---|---|---|---|
| Tian | Sequencing Batch Effect | Pseudo cell mixtures of ADENOCA cell lines | 4 | 577 |
| Kagohara | CTX Treatment | HNSCC Cells | 3 | 23,914 |
| Kang | INF-Beta Treatment | PBMCs | 8 | 24,368 |

**Supplemental Table 5. Summary table of real datasets used to measure performance.**

| Dataset | Method | ARI | LISI | DB |
|---|---|---|---|---|
| Tian | ICAT | 0.65 | 0.30 | 0.79 |
| | No Int. | 0.46 | 0.21 | 0.00 |
| | Seurat 3.1 | 0.69 | 0.27 | 0.42 |
| | Seurat 3.1 + ICAT | 0.67 | 0.32 | 0.83 |
| | Scanorama | 0.65 | 0.17 | 0.22 |
| | Scanorama + ICAT | **0.81** | 0.37 | **1.00** |
| | Harmony | 0.69 | 0.20 | 0.00 |
| | Harmony + ICAT | 0.60 | **0.46** | 0.95 |
| Kagohara | ICAT | **1.00** | 0.65 | 0.95 |
| | No Int. | 0.33 | 0.18 | 0.00 |
| | Seurat 3.1 | 0.54 | 0.61 | 0.10 |
| | Seurat 3.1 + ICAT | **1.00** | 0.61 | 0.55 |
| | Scanorama | **1.00** | 0.35 | 0.50 |
| | Scanorama + ICAT | **1.00** | 0.71 | **1.00** |
| | Harmony | 0.63 | 0.45 | 0.19 |
| | Harmony + ICAT | 0.99 | **0.82** | 0.90 |
| Kang | ICAT | **0.71** | 0.81 | **1.00** |
| | No Int. | 0.32 | 0.53 | 0.00 |
| | Seurat 3.1 | 0.58 | 0.71 | 0.11 |
| | Seurat 3.1 + ICAT | 0.51 | 0.64 | 0.31 |
| | Scanorama | 0.50 | 0.56 | 0.48 |
| | Scanorama + ICAT | 0.57 | 0.75 | 0.65 |
| | Harmony | 0.58 | 0.73 | 0.40 |
| | Harmony + ICAT | 0.63 | **0.91** | 0.00 |

**Supplemental Table 6. Performance in real datasets.**

| ICAT G-Test Results | | | |
|---|---|---|---|
| cluster | pvals | pvals.adj | sig |
| 0 | 0.219566892 | 0.219566892 | FALSE |
| 1 | 0.001212742 | 0.002021236 | TRUE |
| 2 | 0.017265074 | 0.021581343 | TRUE |
| 3 | 4.67E-11 | 1.17E-10 | TRUE |
| 4 | 1.09E-19 | 5.43E-19 | TRUE |

**Supplemental Table 7. Gtest results for differential abundance of ICAT-identified PMC subpopulations between treatments.**

**ICAT Fisher Exact Post-hoc Pairwise Comparisons**

| cluster | odds | pvals | pvals.adj | comparison | sig |
|---|---|---|---|---|---|
| 0 | 2.26127321 | 0.006282516 | 0.012565032 | Chlorate-Control | TRUE |
| 1 | 1.319871795 | 0.415136752 | 0.55351567 | Chlorate-Control | FALSE |
| 2 | 0.301960784 | 0.000140166 | 0.000560666 | Chlorate-Control | TRUE |
| 3 | 0 | 1 | 1 | Chlorate-Control | FALSE |
| 0 | 0.507758621 | 0.047208202 | 0.047208202 | MK886-Control | TRUE |
| 1 | 2.3075 | 0.011937173 | 0.01591623 | MK886-Control | TRUE |
| 2 | 0.011904762 | 1.71E-15 | 6.83E-15 | MK886-Control | TRUE |
| 3 | 0 | 7.94E-14 | 1.59E-13 | MK886-Control | TRUE |

**Supplemental Table 8. Post hoc pairwise Fisher Exact tests for treatment-affected ICAT subpopulations.**

| | Sm50+/Pks2- Model | | | | | |
|---|---|---|---|---|---|---|
| | Coef. | Std.Err. | t | P>ltl | 95% CI Lower | 95% CI Upper |
| Intercept | -2.864186788 | 0.179791482 | -15.93060337 | 3.89E-57 | -3.216571617 | -2.511801959 |
| C(treatment)[T.MK886] | -0.713413802 | 0.216528908 | -3.294773929 | 0.000985009 | -1.137802664 | -0.28902494 |
| C(fert)[T.replicate2] | -0.144183659 | 0.229814507 | -0.627391462 | 0.530402663 | -0.594611815 | 0.306244497 |
| C(fert)[T.replicate3] | -1.606548568 | 0.336857153 | -4.769228005 | 1.85E-06 | -2.266776456 | -0.94632068 |

**Supplemental Table 9. Linear model summary for abundance of *sm50+/pks2-* PMC between DMSO and MK886 treated embryos.**

| | Sm50-/Pks2+ Model | | | | | |
|---|---|---|---|---|---|---|
| | Coef. | Std.Err. | t | P>ltl | 95% CI Lower | 95% CI Upper |
| Intercept | -6.200930225 | 0.605683932 | -10.23789786 | 1.34E-24 | -7.388048917 | -5.013811534 |
| C(treatment)[T.MK886] | 2.603132373 | 0.592924937 | 4.390323651 | 1.13E-05 | 1.441020851 | 3.765243894 |
| C(fert)[T.replicate2] | -0.970285878 | 0.497035923 | -1.952144366 | 0.050921067 | -1.944458387 | 0.003886631 |
| C(fert)[T.replicate3] | 0.417259722 | 0.26968222 | 1.547227409 | 0.121808419 | -0.111307716 | 0.94582716 |

**Supplemental Table 10. Linear model summary for abundance of *sm50-/pks2+* PMC between DMSO and MK886 treated embryos.**

| Seurat G-Test Results | | | |
|---|---|---|---|
| **cluster** | **pvals** | **pvals.adj** | **sig** |
| 0 | 0.86939774 | 0.86939774 | FALSE |
| 1 | 0.770959229 | 0.86939774 | FALSE |
| 2 | 0.300698296 | 0.86939774 | FALSE |
| 3 | 0.454597177 | 0.86939774 | FALSE |
| 4 | 0.253074595 | 0.86939774 | FALSE |
| 5 | 0.64849922 | 0.86939774 | FALSE |

**Supplemental Table 11. Gtest results for differential abundance of Seurat-identified PMC subpopulations between treatments.**

| Scanorama G-Test Results | | | |
|---|---|---|---|
| **cluster** | **pvals** | **pvals.adj** | **sig** |
| 0 | 0.553182475 | 0.553182475 | FALSE |
| 1 | 0.100985069 | 0.168308448 | FALSE |
| 2 | 0.000176536 | 0.000882678 | TRUE |
| 3 | 0.063621448 | 0.159053619 | FALSE |
| 4 | 0.323729775 | 0.404662219 | FALSE |

**Supplemental Table 12. Gtest results for differential abundance of Scanorama-identified PMC subpopulations between treatments.**

| Scanorama Fisher Exact Post-hoc Pairwise Comparisons | | | | | |
|---|---|---|---|---|---|
| **cluster** | **odds** | **pvals** | **pvals.adj** | **comparison** | **sig** |
| 2 | 0.43445122 | 0.00459723 | 0.00459723 | Chlorate-Control | TRUE |
| 2 | 0.219722456 | 1.07E-05 | 2.13E-05 | MK886-Control | TRUE |

**Supplemental Table 13. Post hoc pairwise Fisher Exact tests for treatment-affected Scanorama subpopulations.**

| **Harmony G-test Results** | | | |
|---|---|---|---|
| cluster | pvals | pvals.adj | sig |
| 0 | 0.233265435 | 0.266589069 | FALSE |
| 1 | 0.050488234 | 0.086841879 | FALSE |
| 2 | 0.054276174 | 0.086841879 | FALSE |
| 3 | 0.485554565 | 0.485554565 | FALSE |
| 4 | 0.080876994 | 0.107835992 | FALSE |
| 5 | 1.92155E-06 | 1.53724E-05 | TRUE |
| 6 | 0.019345836 | 0.051588897 | FALSE |
| 7 | 0.004047702 | 0.016190807 | TRUE |

**Supplemental Table 14. Gtest results for differential abundance of Harmony-identified PMC subpopulations between treatments.**

| Harmony Fisher Exact Post-hoc Pairwise Comparisons | | | | | |
|---|---|---|---|---|---|
| **odds** | **pvals** | **pvals.adj** | **cluster** | **comparison** | **sig** |
| 0.591111111 | 0.669645023 | 0.669645023 | 5 | Chlorate-Control | FALSE |
| 2.278911565 | 0.452521258 | 0.603361677 | 7 | Chlorate-Control | FALSE |
| 8.286604361 | 0.000103427 | 0.000413709 | 5 | MK886-Control | TRUE |
| 7.640350877 | 0.002506892 | 0.005013785 | 7 | MK886-Control | TRUE |

**Supplemental Table 15. Post hoc pairwise Fisher Exact tests for treatment-affected Harmony subpopulations.**

| Dataset | alpha | sigma | reg | reference |
|---|---|---|---|---|
| Simulated | 0.1 | 3 | 0.5 | "all" |
| Tian | 0.1 | 3 | 3 | "all" |
| Kagohara | 0.1 | 3 | 3 | "controls" |
| Kang | 0.1 | 3 | 3 | "controls" |
| PMC | 0.1 | 3 | 0.5 | "all" |

**Supplemental Table 16. NCFS hyper-parameters used for each experiment.**

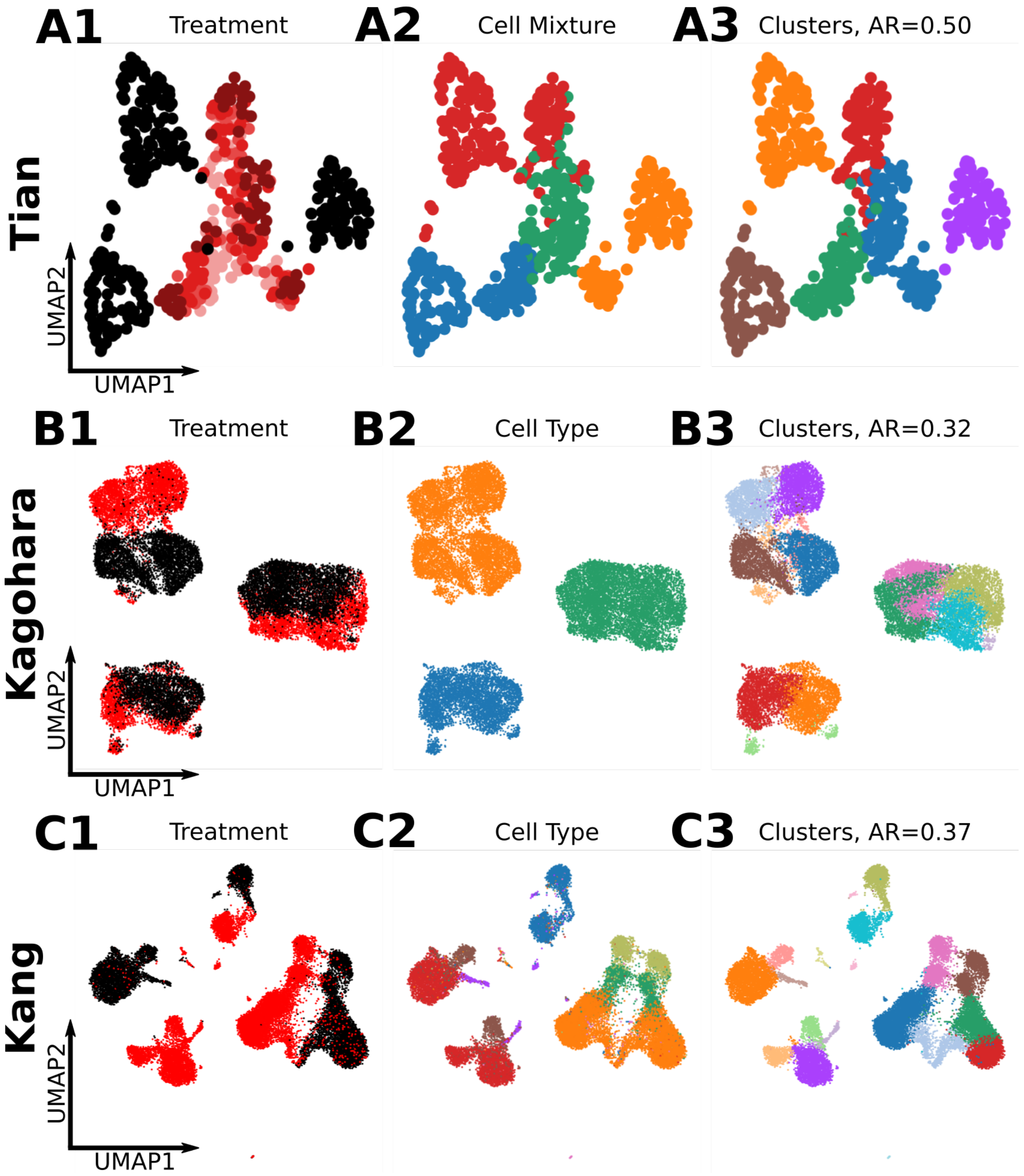


**Supplemental Figure 1. Clustering agnostic to treatment status poorly isolates known cell identities.** UMAPs of cells across three datasets (rows) showing treatment status (column 1), known cell identity (column 2), and identified clusters using the Louvain method (column 3) are shown for Tian (A), Kagohara (B), and Kang (C) datasets. Treatment UMAPs (left) display control cells as black dots, while red dots represent treated cells. For cell identity and cluster UMAPs (middle and right), each cell is colored by cell identity or cluster, respectively. The mismatch between produced cluster labels and known cell identities demonstrates a failure to recapitulate known populations.dataset. AR; adjusted rand index calculated from known cell identities and produced cluster labels.

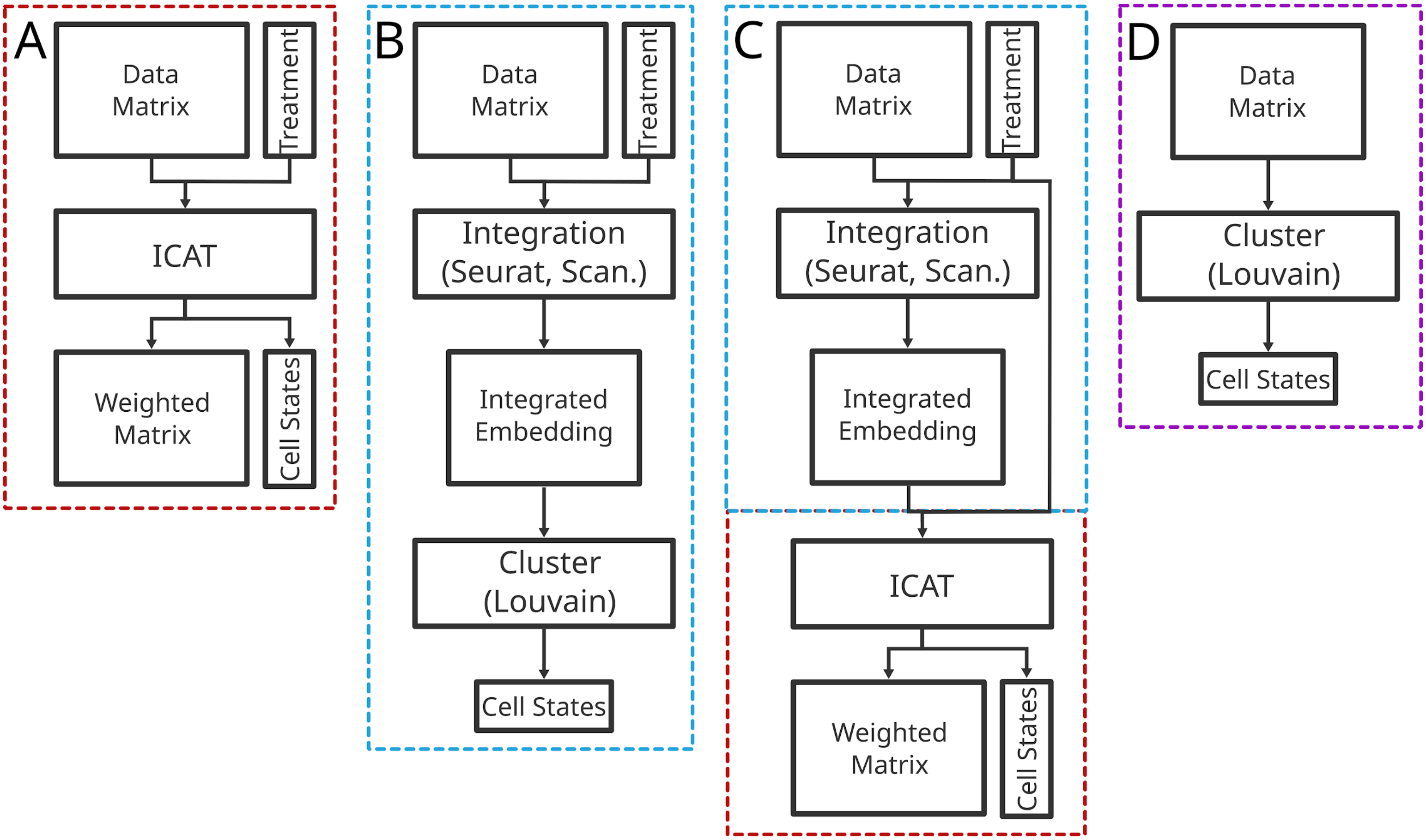


**Supplemental Figure 2. Schematic of workflow comparisons for cell state identification in perturbation experiments.** Cell states are identified by one of four methods: **A.** The pre-processed count matrix and treatment status is provided to ICAT for cell-state identification. **B.** Treatment-separated count matrices are integrated using an integration algorithm (Seurat or Scanorama) to produce a joint embedding. Cells are clustered in the joint embedding using a traditional clustering algorithm (Louvain). **C.** Treatment-separated count matrices are integrated using an integration algorithm (Seurat or Scanorama) to produce a joint embedding. The embedded matrix and treatment status are provided to ICAT for cell-state identification. **D.** No consideration for treatment is provided, and cells are clustered naively using a traditional clustering algorithm (Louvain).

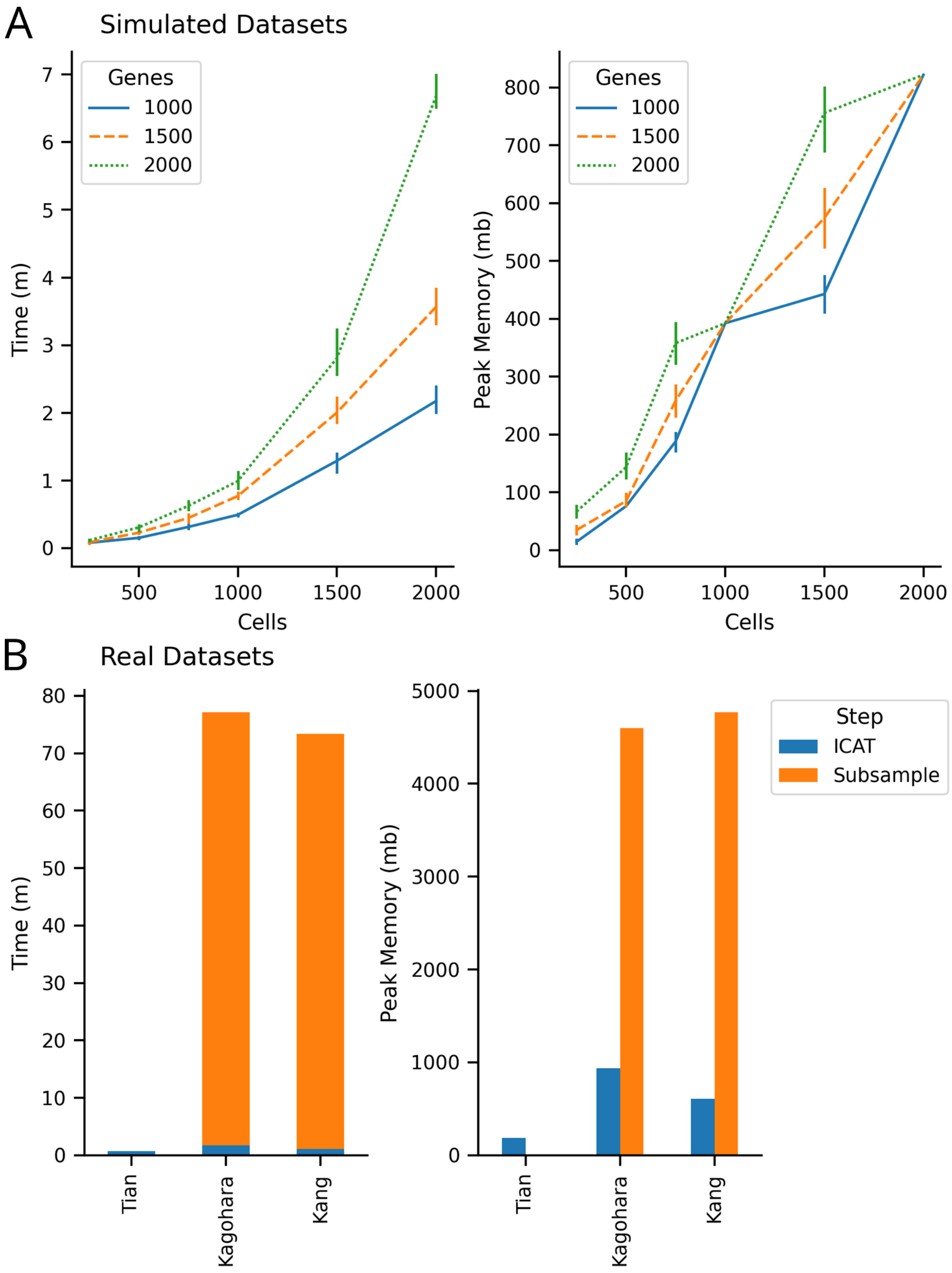


**Supplemental Figure 3. Timing and memory performance of ICAT.** Time to convergence and memory consumption of ICAT over simulated datasets (A) and real datasets (B). **A.** Simulated datasets were simulated with varying numbers of cells (x-axis) and genes (color). Left panel shows complete run time, while the right shows memory consumption. Each cell-gene combination was simulated three times (n = 3). Error bars represent the 95% confidence interval for mean values. **B.** Stepwise breakdown of time to convergence and memory consumption in real datasets for submodular optimization subsampling (orange) and ICAT (blue). Area represents total time (left panel) or peak memory usage (right panel). Number of cells for each dataset: Tian (n = 577), Kagohara (n = 23,914), Kang (n = 24,368). No subsampling was performed for the Tian dataset.

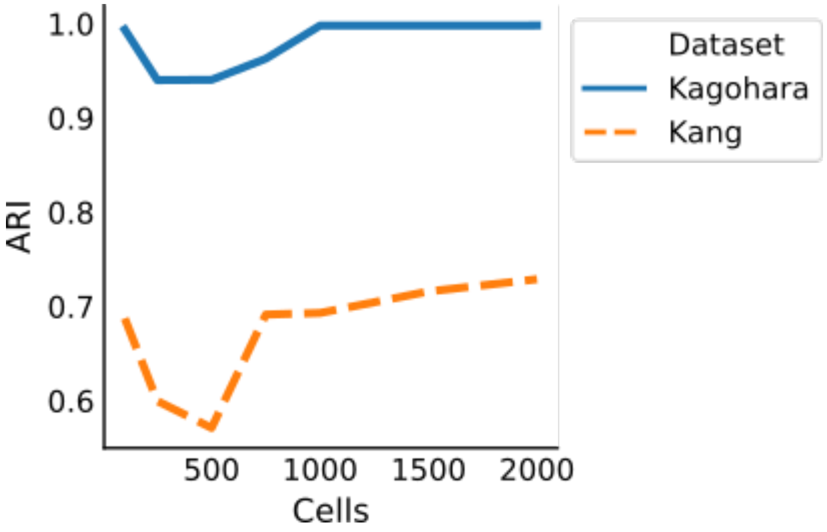


**Supplemental Figure 4. ICAT efficiently accounts for the heterogeneity in datasets using a small number of cells.** The performance of ICAT was measured using ARI for the Kang and Kagohara datasets as the number of cells used for NCFS feature weighting is increased. The number of cells to use ranged from 100 to 2000. Representatives cells were selected via submodular optimization using the FacilitySelector in *Apricot*. Both datasets show strong performance for low cell numbers (n=100), with a dip at intermediate values (n=250, n=500), and tapered performance gain for higher number (n=750-2000), indicating only a small number of cells are necessary to efficiently learn the data space.

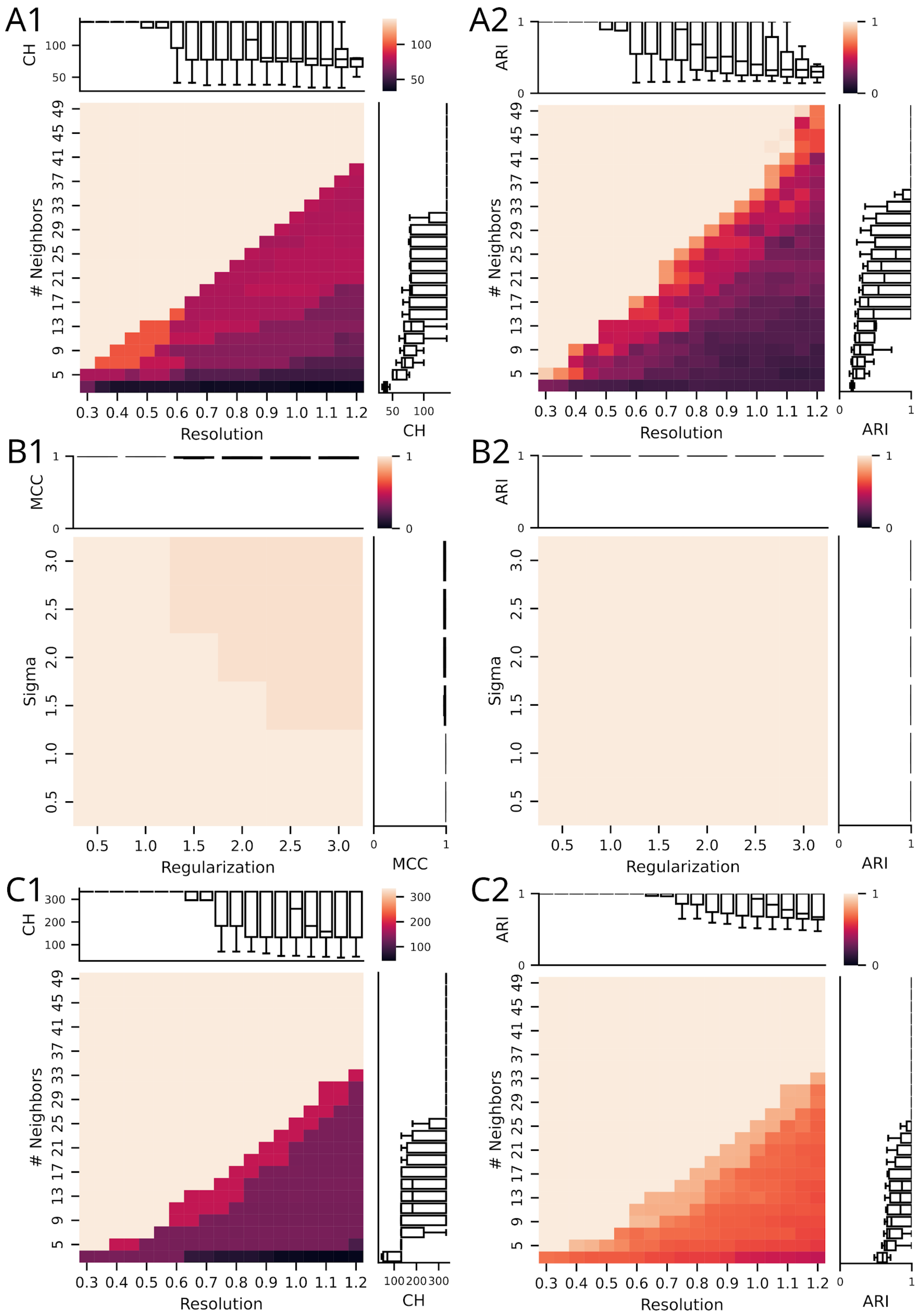


**Supplemental Figure 5. Grid search correctly tunes ICAT hyper parameters.**
Performance of ICAT over varying hyper parameter combinations for each step in the algorithm: identifying control cell states via Louvain (A), weighting highly predictive genes via NCFS (B), and the final Louvain clustering with immutable control states (C). Parameters were optimized for each sequential step, with default parameters being used for any downstream steps. **A.** Performance of ICAT when altering the number of neighbors during the KNN graph construction (rows) and the resolution parameter during Louvain (columns). Color represents either the CH criterion (A1), which is maximized during grid search, or the ARI measuring global label agreement between ground truth labels and final labels produced by ICAT (A2). Boxplots show marginal distributions for each parameter. **B.** Performance of ICAT when altering the kernel width parameter (sigma, rows) and regularization parameter (columns) during NCFS. Color represents either the MCC over a three-fold cross validation (B1), which is maximized during grid search, or the ARI. **C.** Performance of ICAT when altering the number of neighbors during the KNN graph construction (rows) and the

resolution parameter during Louvain (columns). Color represents either the CH score (A1), which is maximized during grid search, or the ARI measuring global label agreement between ground truth labels and final labels produced by ICAT (A2). Boxplots show marginal distributions for each parameter. KNN, k-nearest neighbors; CH, Calinski-Harabasz criterion; ARI, Adjusted Rand Index; MCC, Matthew's Correlation Coefficient.

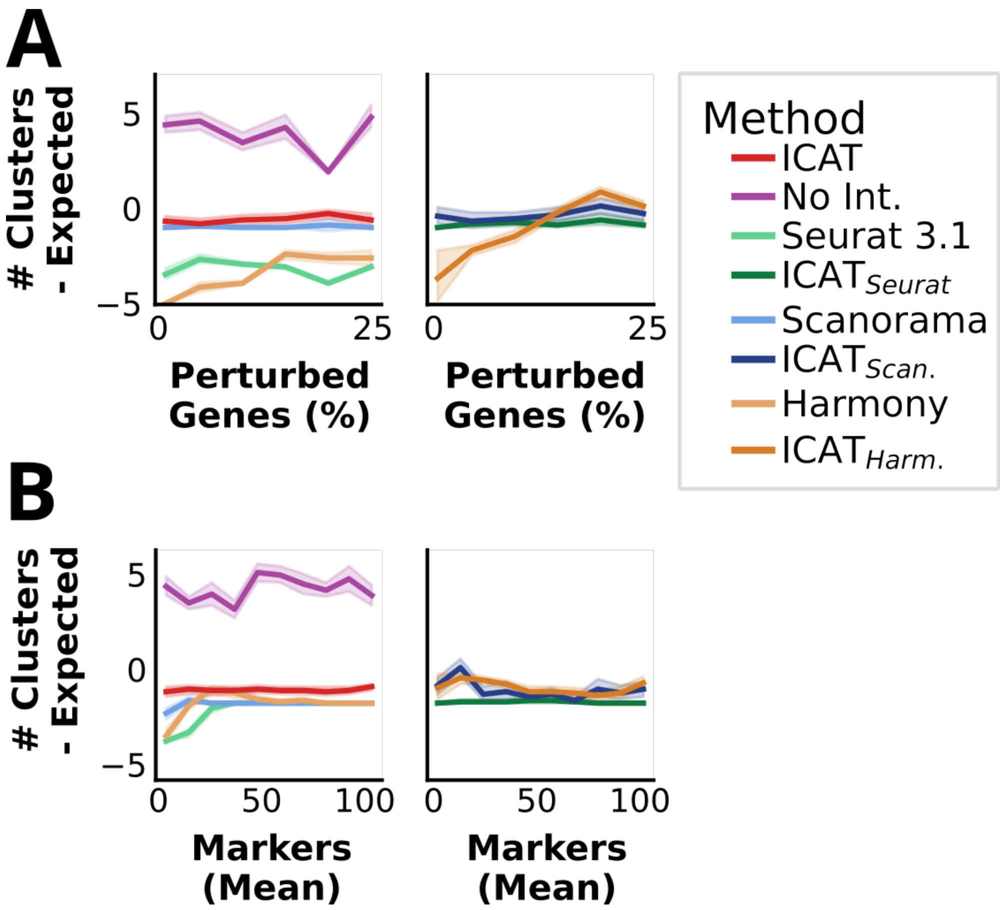


**Supplemental Figure 6. ICAT identifies the correct number of cell states across a range of perturbation and signal intensities.** Each plot compares the performance of the indicated algorithms on simulated data, displaying the deviation from expected number of clusters (y-axis) as the percent of perturbed genes (A) or the number of marker genes per cell identity (B) increases. Negative values indicate under clustering (fewer observed clusters compared to known labels) while positive values indicate over clustering. **A.** Comparison between ICAT, Seurat, Scanorama, Harmony, and no integration (left) as perturbation severity increases. No integration severely over clusters cells while Seurat and Harmony generally under clusters.I ICAT$_{Seurat}$ and ICAT$_{Harm.}$(right) generally rescues under clustering found in Seurat and Harmony, respectively. **B.** Comparison between ICAT, Seurat, Scanorama, Harmony, and no integration (left) as the number of marker genes per cell state increases. ICAT$_{Seurat}$ and ICAT$_{Harm.}$(right) generally rescues under clustering found in Seurat and Harmony at lower signal resolutions, respectively.

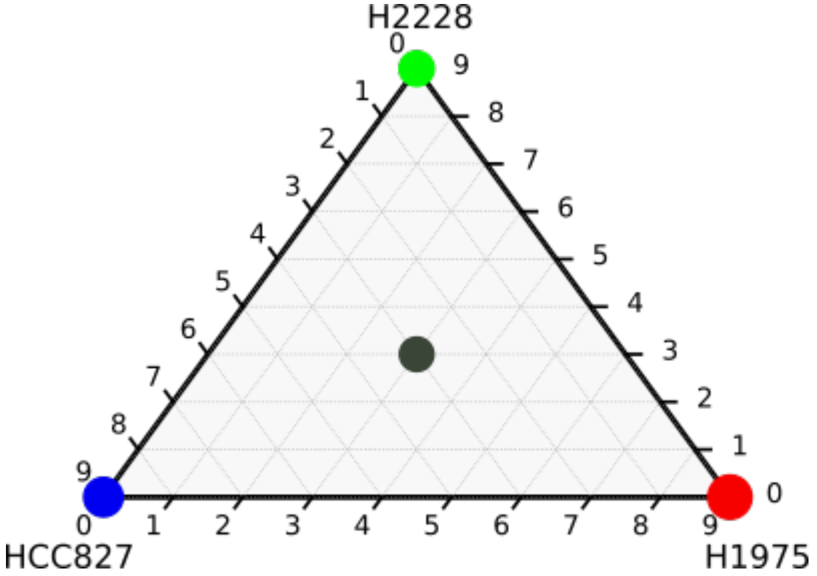


**Supplemental Figure 7. The Tian dataset is composed of four cell types.** The pure cells and 1:1:1: cell mixture that were used herein to evaluate the performance of ICAT. The dots are colored by mixture of each cell line, and the size of each dot is proportional to the number of cells in each cell mixture. Adapted from Tian et al.[21]

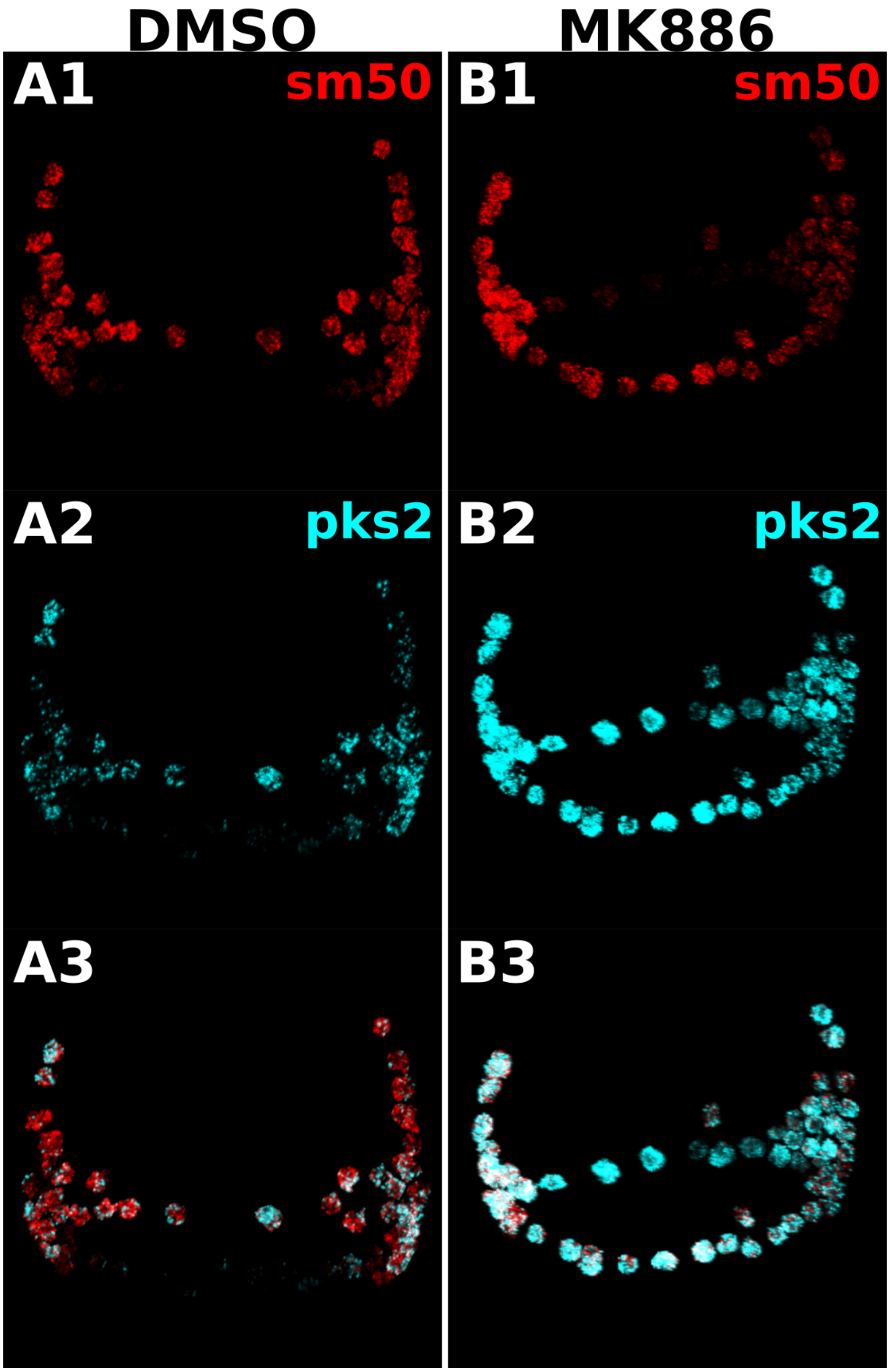


**Supplemental Figure 8. MK886-treated embryos display distinct *sm50* and *pks2* expression patterns compared to control.** HCR FISH images from Fig. 4B for *sm50* (red) and *pks2* (cyan) in DMSO (A) and MK886-treated (B) embryos are shown individually (1, 2) and merged (3).

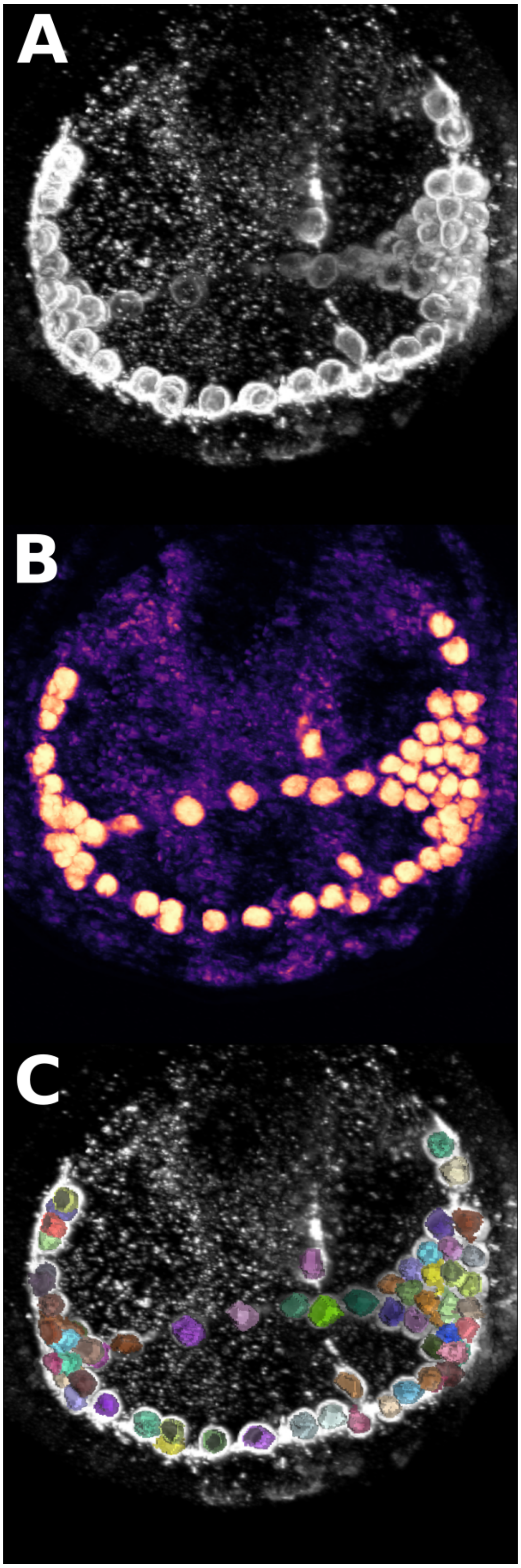


**Supplemental Figure 9. A random forest classifier correctly identifies and segments PMCs in sea urchin embryos. A.** Raw PMC stain used to predict PMC probabilities for each voxel in confocal images. **B.** PMC probabilities generated by Random Forest classifier. Brighter colors (yellow / reds) represent higher probabilities, while darker colors (blacks / purple) represent low probabilities. **C.** Final PMC segmentation produced by post processing of PMC probabilities.

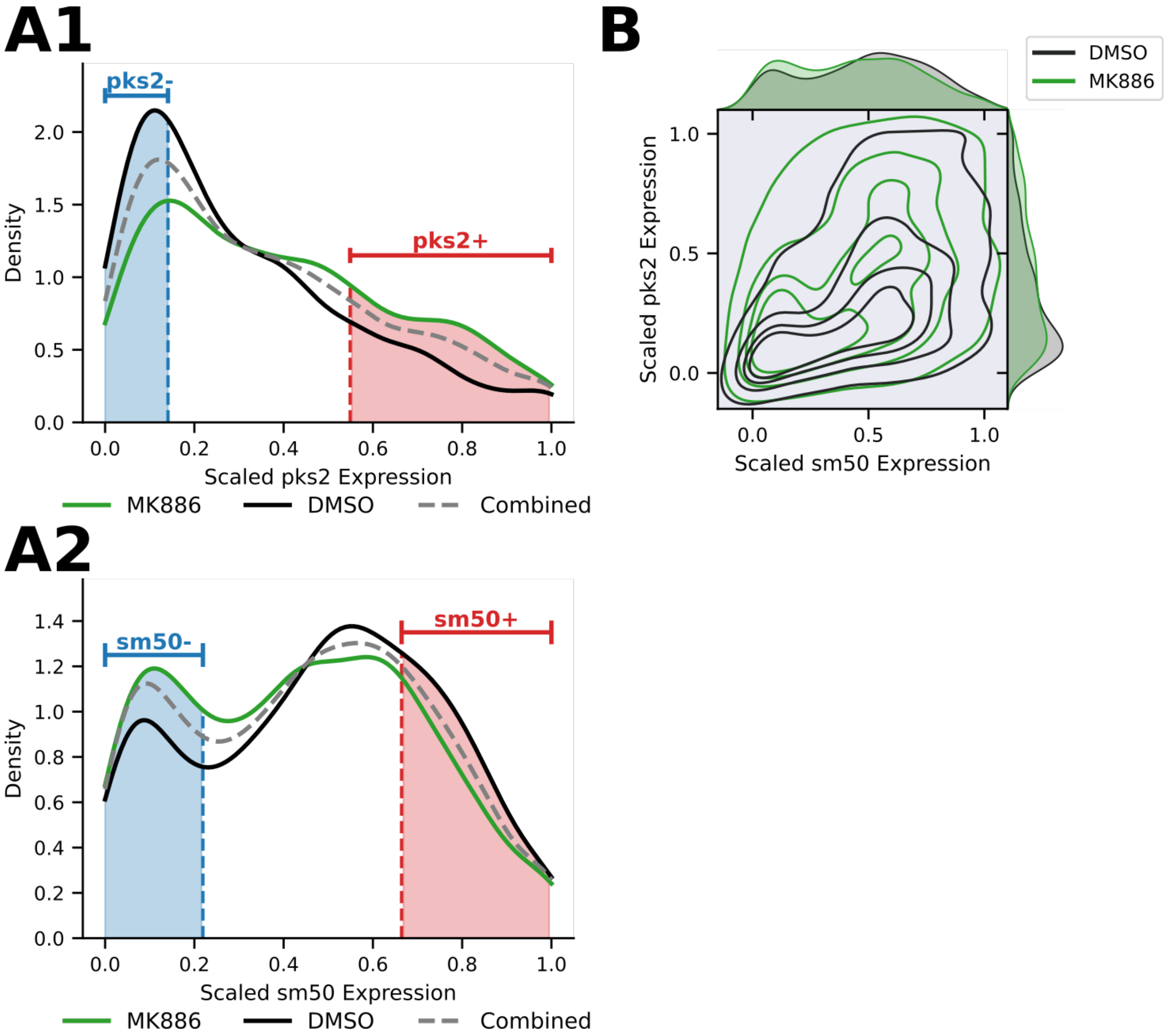


**Supplemental Figure 10. Definition of under and over-expression of *pks2* and *sm50*. A.** Distribution of scaled pks2 (A1) and sm50 (A2) expression in cells from HCR FISH data. The $25^{th}$ and $75^{th}$ percentiles (shaded) were selected for threshold to determine under (-) and over (+) gene expression, respectively. **B.** Joint distribution of pks2 and sm50 HCR FISH expression profiles.

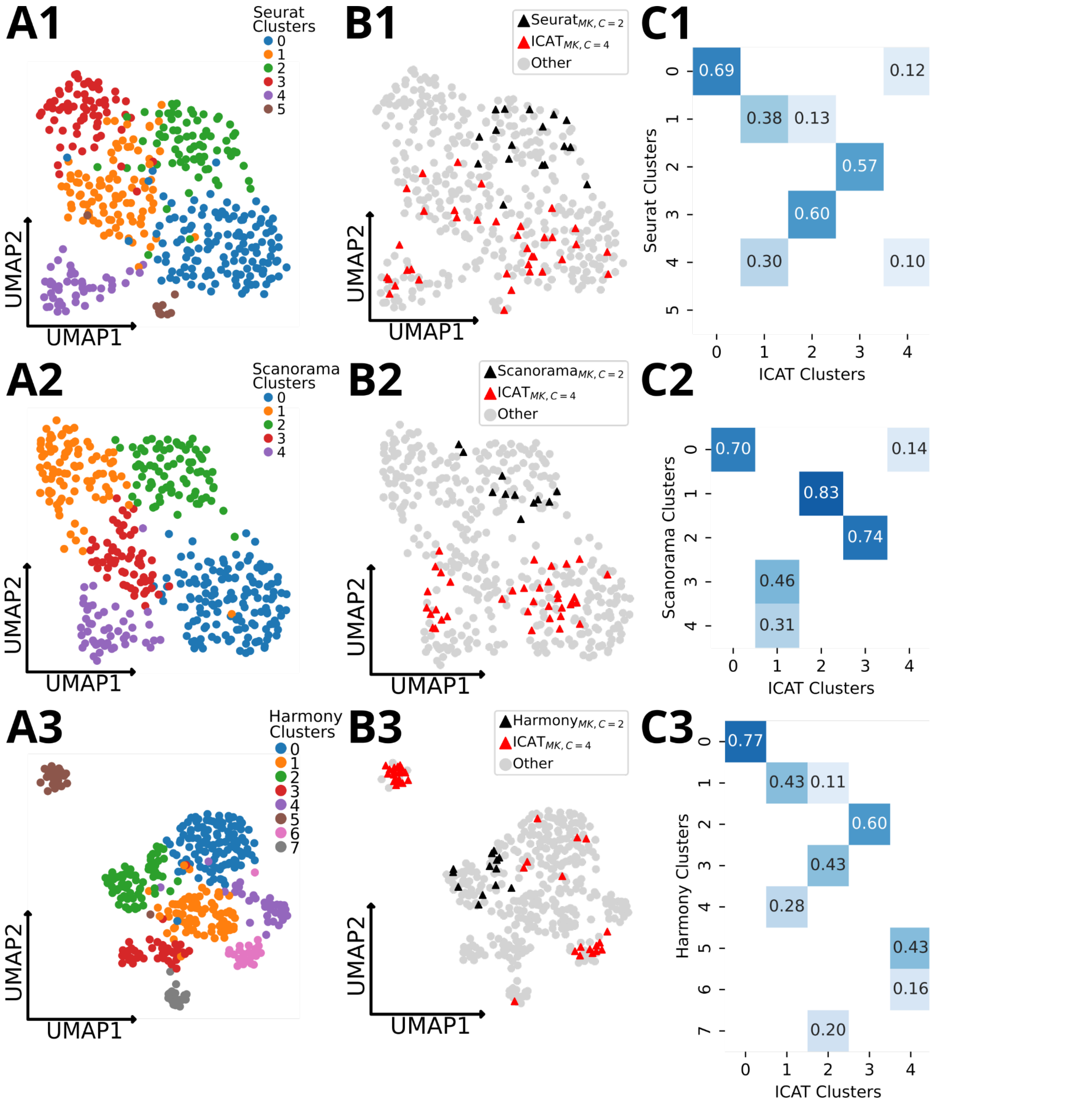


**Supplemental Figure 11. Integration methods fail to correctly define the response of PMCs to MK886 treatment. A.** UMAP projections display clusters generated by Seurat (A1), Scanorama (A2), and Harmony (A3) workflows. **B.** UMAP projections highlighting either MK-treated cells expected to be absent from the closest sm50+/pks2- analogue detected by integration workflows (black) or ICAT-identified sm50-/pks2+ MK-treated cells (red). Seurat (B1) includes many MK cells in its expected sm50+/pks2- analogue, while distributing sm50-/pks2+ MK cells between multiple clusters. Scanorama (B2) shows some absence of MK cells in the analogous sm50+/pks2- cluster, but dispersal of the reciprocal sm50-/pks2+ state between three clusters. Harmony (B3) correctly isolates the MK-induced sm50-/pks2+ PMC cells state, while still incorporating many MK cells in its sm50+/pks2+ cluster analogue. **C.** Jaccard similarity between ICAT-identified clusters and those produced by each integration workflow (Seurat C1, Scanorama C2, Harmony C3). Scores < 0.1 are masked. Cell state analogues in (B) were matched via the highest jaccard score for ICAT cluster 3.

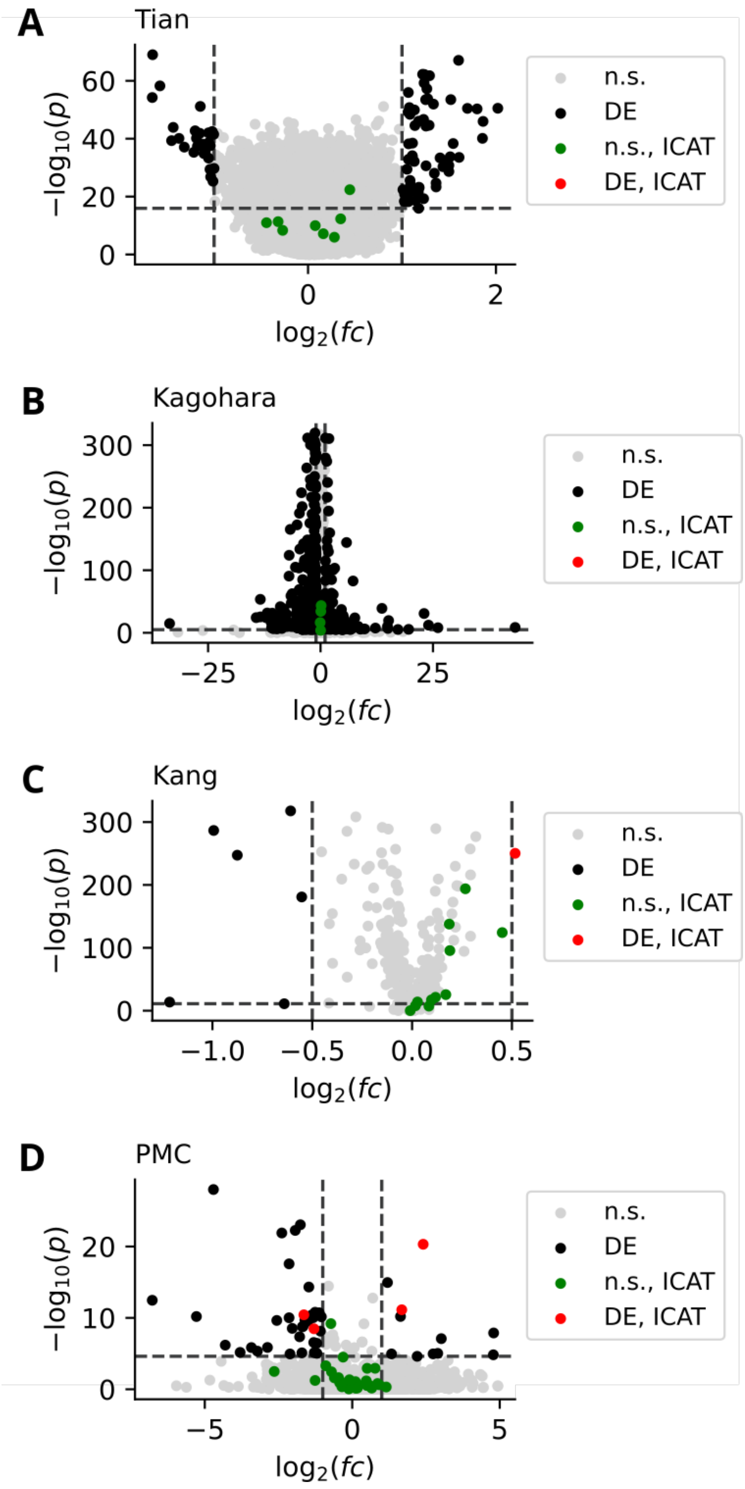


**Supplemental Figure 12: ICAT-selected cell state marker genes are largely stable between treatments.** Volcano plots showing differentially expressed genes between treatments in the Tian (A), Kagohara (B), Kang (C) and PMC SMARTSeq (D) datasets. The Y-axis shows the -log10 nominal p-value while X-axis shows the log2 fold change in gene expression compared to controls. Genes were considered differentially expressed with fdr < 0.05 and | log2(fc) | > 1 (represented by dashed lines) except for the Kang dataset, where the fold change threshold was set to log2(1.5). Green and red dots represent informative genes identified by ICAT (weight > 1), where genes colored green were not found to be differentially expressed while red were.
